# Multi-Targeted Effects of Novel Cycloastragenol Derivatives: Enhancing NRF2, Proteostasis, and Telomerase Pathways with p53 Modulation to Delay Replicative Senescence

**DOI:** 10.1101/2024.08.12.607434

**Authors:** Sinem Yilmaz, Erdal Bedir, Petek Ballar Kirmizibayrak

## Abstract

Aging is a complex, multifactorial process driven by various cellular and molecular mechanisms, including telomere shortening, oxidative stress, and the decline of proteostasis, all of which contribute to replicative senescence and age-related diseases. Cycloastragenol (**CA**), a triterpenoid saponin derived from *Astragalus membranaceus*, has shown potential for its ability to activate telomerase, suggesting therapeutic benefits in delaying cellular aging. In this study, we explored the effects of novel **CA** derivatives, obtained through biotransformation as telomerase activators, on the NRF2/proteasome/telomerase axis and their potential to delay replicative senescence in human primary epidermal keratinocytes (HEKn).

Our findings reveal that these **CA** derivatives significantly enhance NRF2 nuclear activity, leading to the upregulation of key cytoprotective enzymes essential for mitigating oxidative stress. Notably, these derivatives exhibited efficacy at much lower concentrations compared to **CA**, demonstrating their potential for enhanced therapeutic application. The derivatives also markedly increased proteasome activity, particularly in the β1 and β5 subunits, thereby preserving proteostasis—a critical factor in preventing the accumulation of damaged proteins associated with aging. Furthermore, continuous treatment with these derivatives sustained stimulatory effects, which was evidenced by increased NRF2, proteasome, and hTERT protein levels even in senescent cells and extended cellular lifespan.

Additionally, we explored the impact of **CA** derivatives on p53-mediated pathways, demonstrating that these compounds effectively modulate the p53/p21 axis, reducing cell cycle arrest and promoting cellular proliferation. Moreover, the derivatives exhibited neuroprotective properties by attenuating glutamate-induced excitotoxicity, further underscoring their potential as multi-targeted anti-aging agents. In conclusion, our study provides strong evidence that novel **CA** derivatives act on multiple fronts to enhance NRF2 activity, maintain proteostasis, and modulate telomerase and p53 pathways, most at lower doses compared to **CA**. These actions collectively contribute to the delay of replicative senescence and the promotion of cellular longevity, positioning **CA** derivatives as potent candidates for developing multi-targeted anti-aging therapies that address the complex interplay of aging-related cellular processes.

**Highlights:**

- Telomerase-active **CA** derivatives enhance NRF2 activity and proteasome activity, leading to cytoprotection at lower doses than **CA**.
- **CA** derivatives modulate the p53 pathway and cell cycle, prolonging cellular lifespan and delaying replicative senescence.
- **CA** derivatives protect cells against glutamate-excitotoxicity along with decreased p53 protein levels.

## 1. Introduction

Aging is a complex biological process characterized by a gradual decline in cellular and tissue function, which increases the vulnerability to several diseases (1). This process is driven by a combination of interconnected factors, including telomere shortening, oxidative stress, and proteostasis impairment, all also contributing to the onset of age-related conditions such as cardiovascular and neurodegenerative diseases (2, 3). Telomere shortening limits the replicative capacity of cells, leading to replicative senescence, a significant contributor to the aging phenotype (2). This process triggers the DNA Damage Response, resulting in cell cycle arrest through pathways, which involves key proteins like ATM, p53, and cyclin-dependent kinase (CDK) inhibitors such as p21 and p16 (4–6). Telomerase, comprised of telomerase reverse transcriptase subunit (TERT) and RNA component (TERC), maintains telomere length and genomic stability and delays senescence and cellular aging (5, 7). Similarly, NRF2, the central transcription factor for maintaining redox balance and signal transduction, is another essential regulator of longevity (8, 9). Induction of NRF2-mediated antioxidant defense system is crucial in preventing cellular senescence by simultaneously regulating oxidative stress, endoplasmic reticulum stress, inflammation, and cell cycle arrest (10). However, its expression and function can be suppressed by p53 during senescence (10, 11), which contributes to the decline in the proteostasis network that is essential for maintaining protein homeostasis and cellular function (12) and is closely linked to organismal aging and replicative senescence. Additionally, the proteasome inhibition-mediated senescence has been suggested to be regulated by the p53 pathway (13).

Extensive studies have demonstrated that numerous factors contribute to each aging pathway, and targeting these pathways could delay tissue dysfunction and the onset of age-related diseases (14). Given these interconnected mechanisms, there is a growing interest in identifying novel bioactive molecules with multi-target effects to slow aging and promote a healthier lifespan (15). One such promising molecule is Cycloastragenol (CA), a small molecule telomerase activator known commercially as TA-65®. **CA**, the hydrolysis product of one of the main active components, Astragaloside IV, isolated from *Astragalus* species, exhibits several pharmacological activities, including anti-inflammatory, anti-fibrosis, anti-viral, liver protection, endothelial protection, neuroprotection, and immune regulation (16–20). Studies have demonstrated that **CA** may play a role in regulating some main cellular signaling pathways, such as ERK/JNK, Wnt/β-catenin, AKT1-mTOR-RPS6KB1 and JAK/STAT3 signaling (21–24). Despite several reports indicating that **CA** is a promising natural telomerase activator for treating age-related diseases and promoting healthy aging, the underlying mechanisms have yet to be fully elucidated. Building upon these reports, our recent studies have elucidated the status of **CA** at the interconnection of redox balance, telomerase, and proteasome status and reported that **CA** enhances proteasome activity/assembly, dependent on telomerase activation induced via NRF2, suggesting that **CA** triggers crosstalk at the intersection of these three major cellular pathways for aging (25). Since **CA**-based molecule libraries are the focus of several research groups as well as healthcare and pharmaceutical industries, we conducted microbial transformation studies on **CA** and its production artifact **AG** using endophytic fungi (*Alternaria eureka* 1E1BL1 and *Camarosporium laburnicola* 1E4BL1) and identified novel **CA** and **AG** derivatives by biotransformation, a robust methodology for structural modification of complex molecules to obtain unique compounds with diverse chemistry. These derivatives were initially screened for telomerase activation, six of which were identified as novel potent telomerase activators enhancing hTERT protein levels with higher activity than the lead compound **CA** (26–28) (**Figure 1, Supplementary 1**).

**Figure 1.**
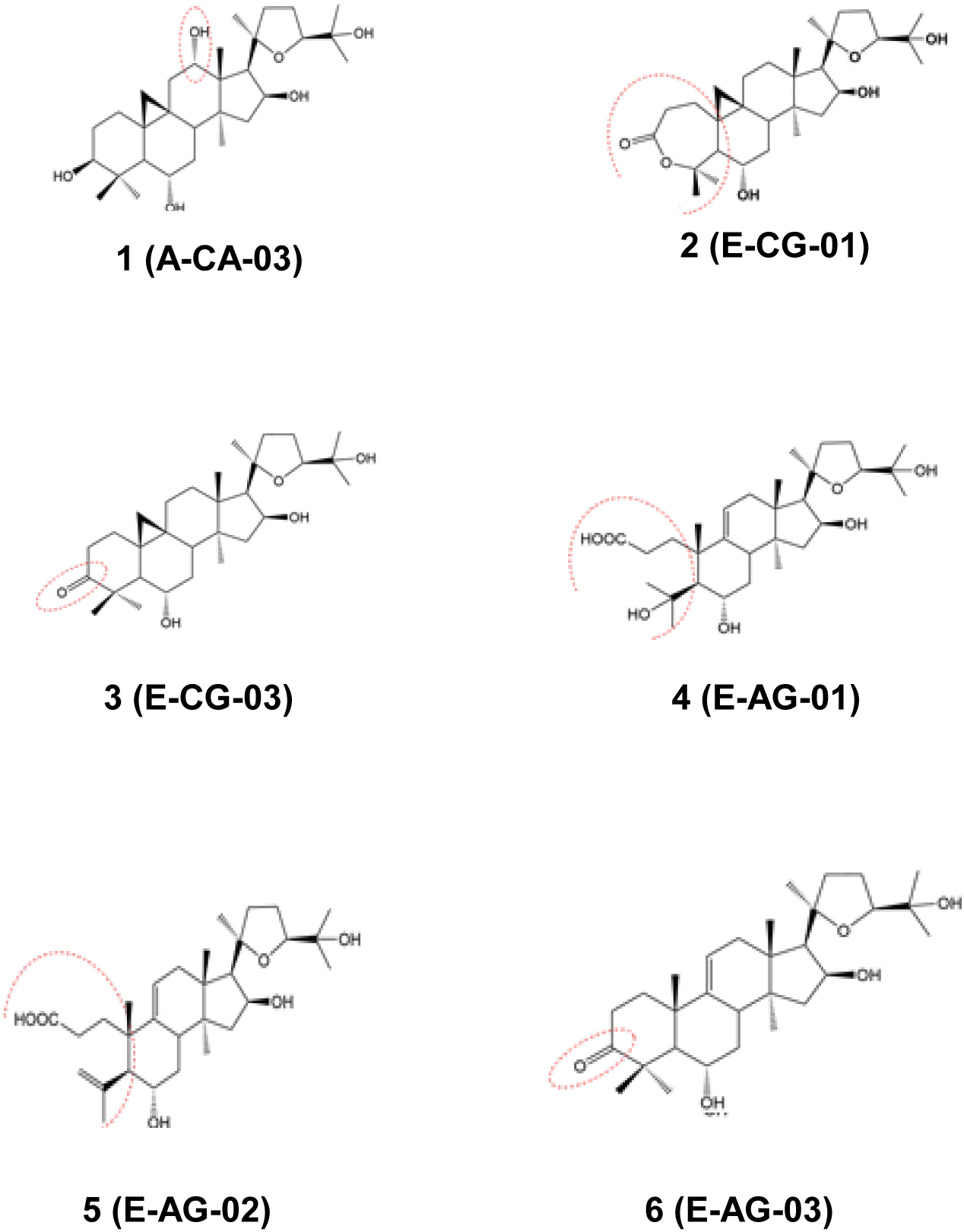
Chemical structure of novel Cycloastragenol derivatives obtained from biotransformation.

Given that telomerase, proteasome, and NRF2 pathways act together to prevent irreversible growth arrest during senescence, and our lead compound CA intersects with these activities, we hypothesized that these derivatives might activate both NRF2 and proteasome and delay replicative senescence via the NRF2/proteasome axis regulated by p53. Considering the interconnected roles of these key players in regulating cellular senescence and proteostasis, targeting them simultaneously could offer a comprehensive strategy to delay aging and promote cellular longevity. This study investigates the potential of six novel **CA** derivatives to activate NRF2, enhance proteasome activity, and modulate the p53 pathway to delay replicative senescence.

## 2. Results

### 2.1. Cytoprotective Efficacy of Novel Cycloastragenol Derivatives through Enhanced NRF2 Activation in Human Epidermal Keratinocytes

The impact of the new derivatives on the NRF2/ARE system was first assessed by measuring the nuclear transcriptional activity of NRF2 in both young (P5) and old (P15) neonatal human epidermal keratinocytes (HEKn). Nuclear NRF2 activity was significantly increased by derivatives **1** and **3** at low nanomolar levels (0.1 and 0.5 nM) and by derivatives **2** and **5** at slightly higher concentrations (2-10 nM and 10-100 nM, respectively) in young HEKn cells (**Figure 2A**). Derivatives **4** and **6** exhibited a concentration-dependent increase in activity at higher concentrations (100 and 1000 nM) (**Figure 2A**). Considering the decline in NRF2 activation during aging (29, 30), the effects of the biotransformation products on P15 old cell HEKn cells were also assessed, revealing that all derivatives enhanced nuclear NRF2 transcriptional activity (**Supplementary S2A).**

**Figure 2.**
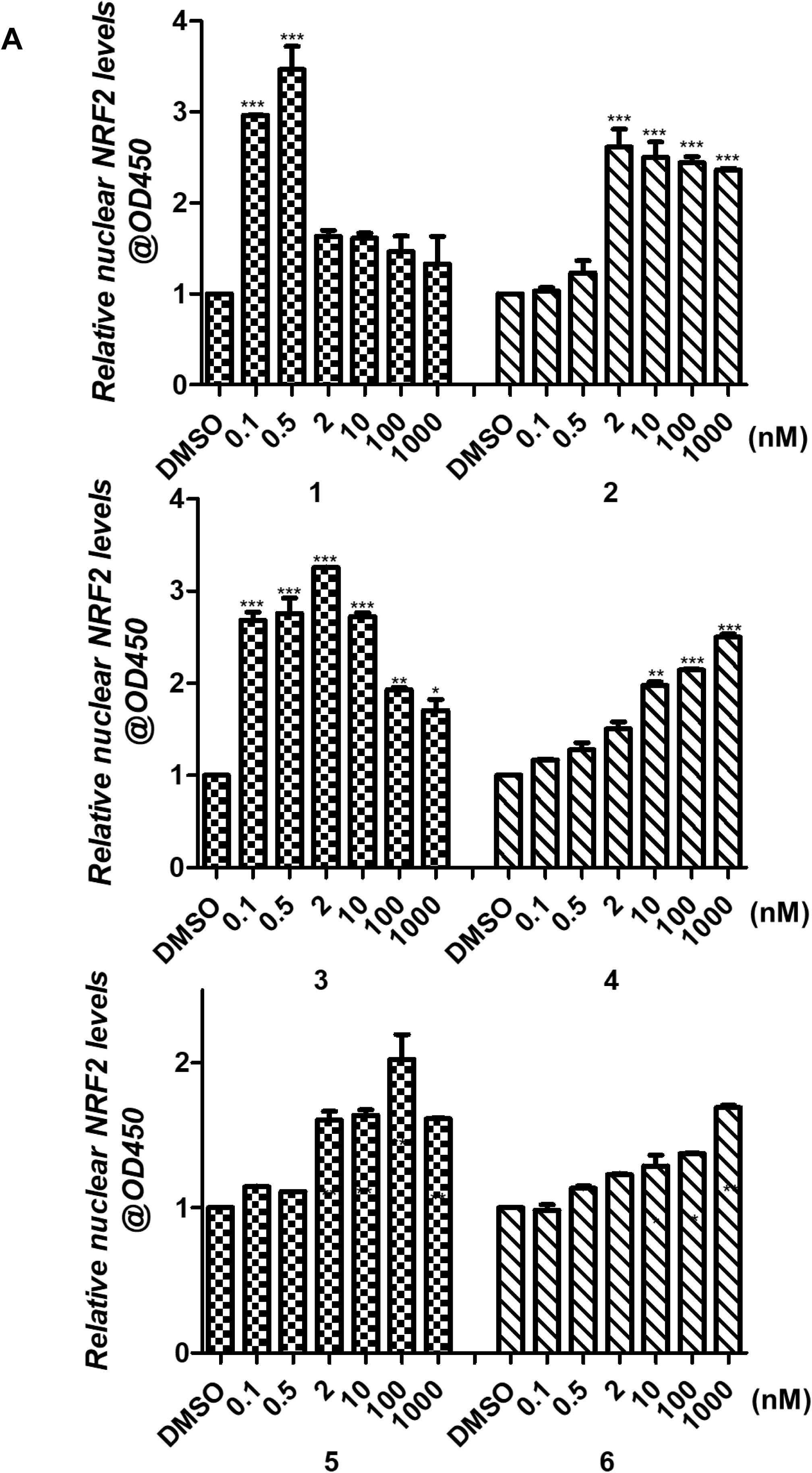
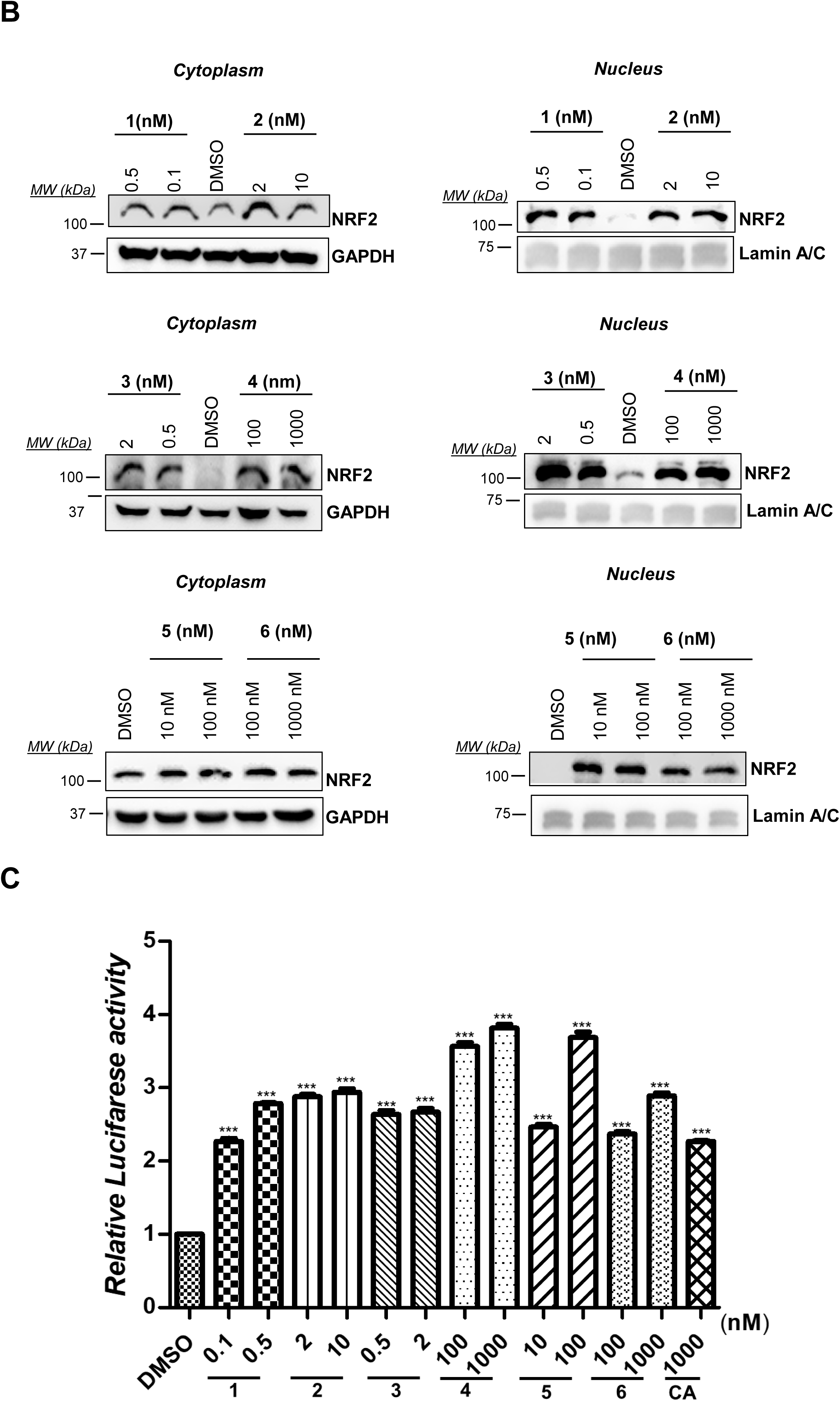
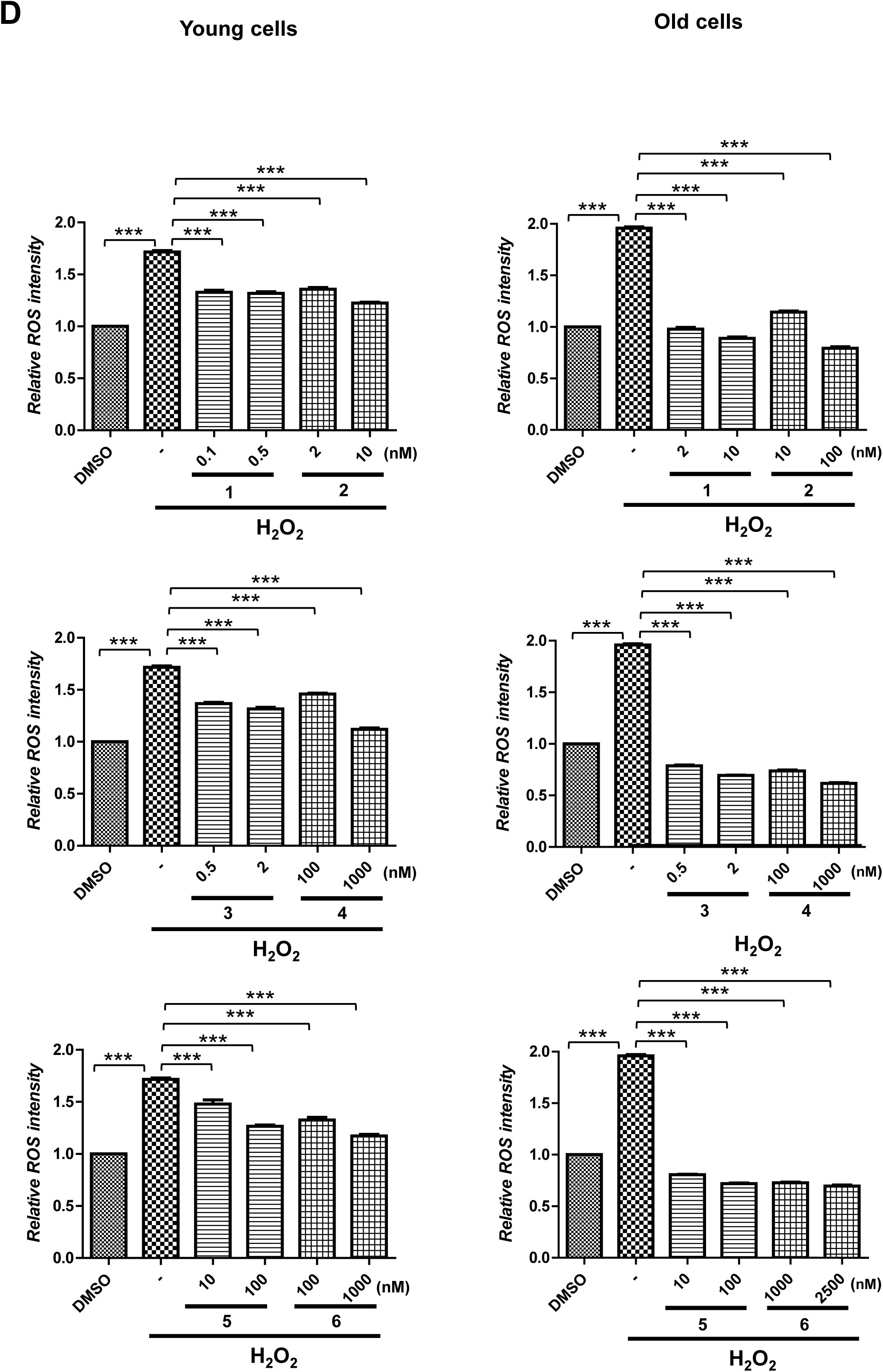
Evaluation of the effects of CA derivatives on the NRF2/ARE system. HEKn cells were treated with an increased concentration of **CA** derivatives for 24 h. (**A**) After nuclear-cytoplasmic fractionation, nuclear lysates were used for NRF2 transcription factor assay with an ELISA kit. After the data obtained from nuclear transcriptional activity was normalized to nuclear protein levels, these activities were presented as fold change compared to the DMSO-treated control cells. Error bars are presented as standard deviations (n = 3; *p ≤ 0.05, **p ≤ 0.001, ***p ≤ 0.005). (**B**) After nuclear-cytoplasmic fractionation, nuclear NRF2 localization was examined by IB. While GAPDH was used for cytoplasmic loading control, Lamin A-C was used for nuclear loading control. (**C**) To determine ARE-promoter activity, a Luciferase activity assay was used. **CA** was used as an experimental control. Error bars are presented as standard deviations (n = 3; *p ≤ 0.05, **p ≤ 0.001, ***p ≤ 0.005). Error bars are presented as standard deviations (n = 3; *p ≤ 0.05, **p ≤ 0.001, ***p ≤ 0.005). (**D**) The ROS scavenging properties of derivatives were assessed both in young and old HEKn cells by H2DCFDA. Error bars are given as standard deviations (n = 3; *p ≤ 0.05, **p ≤ 0.001, ***p ≤ 0.005).

Following the assessment of nuclear NRF2 activity, we examined the effects of these derivatives on NRF2 nuclear localization, ARE promoter activity, NRF2 mRNA expression, and the levels of downstream proteins. Our analysis showed that treatment with derivatives led to enhanced nuclear localization of NRF2 from the cytoplasm significantly increased ARE promoter activity, and upregulated NRF2 mRNA levels (**Figure 2B, 2C, Supplementary S2B- C**). Notably, there was a significant increase in the protein levels of NRF2 and c-JUN, one of the essential leucine zipper region proteins found to be associated with NRF2 to regulate the antioxidant response (31) (**Supplementary S2D**). Further analysis in both young and old HEKn cells revealed that all derivatives markedly enhanced the levels of cytoprotective enzymes such as GR, HO-1, and GCLC, which are transcriptionally regulated by the NRF2/ARE system (21) (**Supplementary S2E)**. In line with these findings, the effects of all derivatives on H_2_O_2_-induced reactive oxygen species (ROS) formation were evaluated in young HEKn cells. We found that all derivatives attenuated ROS formation (**Figure 2D**). As ROS levels are increased during cellular senescence due to the diminished antioxidant response (2), the effect of derivatives on H_2_O_2_-induced ROS formation in P15 old cells was also examined, revealing that the derivatives diminished ROS levels in senescent cells to the levels observed in DMSO-treated control cells (**Figure 2D).**

Taken together, our results indicate that all derivatives enhance NRF2 activation, which in turn upregulates cytoprotective enzymes, conferring protection against oxidative stress induced by H_2_O_2_. These derivatives show promising NRF2 activation at lower concentrations than those previously reported for **CA** (25), suggesting that some derivatives may possess superior bioavailability compared to the lead compound, **CA**, which has poor bioavailability (32).

### 2.2. Proteasomal Activity Enhancement by Novel Cycloastragenol Derivatives

Previous studies have indicated that both NRF2 and telomerase subunit TERT might directly or indirectly regulate proteasomal activity. Thus, the ability of **CA** derivatives to enhance proteasomal function was investigated using specific-fluorogenic substrates for each subunit. Consequently, cells were treated with the derivatives for 24 h to assess their effect on proteasomes. All derivatives increased the activity of β1 (caspase-like; C-L) and β5 (chymotrypsin-like; CT-L) subunits, but not β2 (trypsin-like; T-L) subunit activity (**Figure 3A**). Consistent with these results, all derivatives increased the protein levels of β1 and β5 but not β2 (**Figure 3B**). Additionally, all derivatives also augmented the proteasomal activity in old HEKn cells (**Supplementary S3**), suggesting these **CA** derivatives are novel proteasome activators.

**Figure 3.**
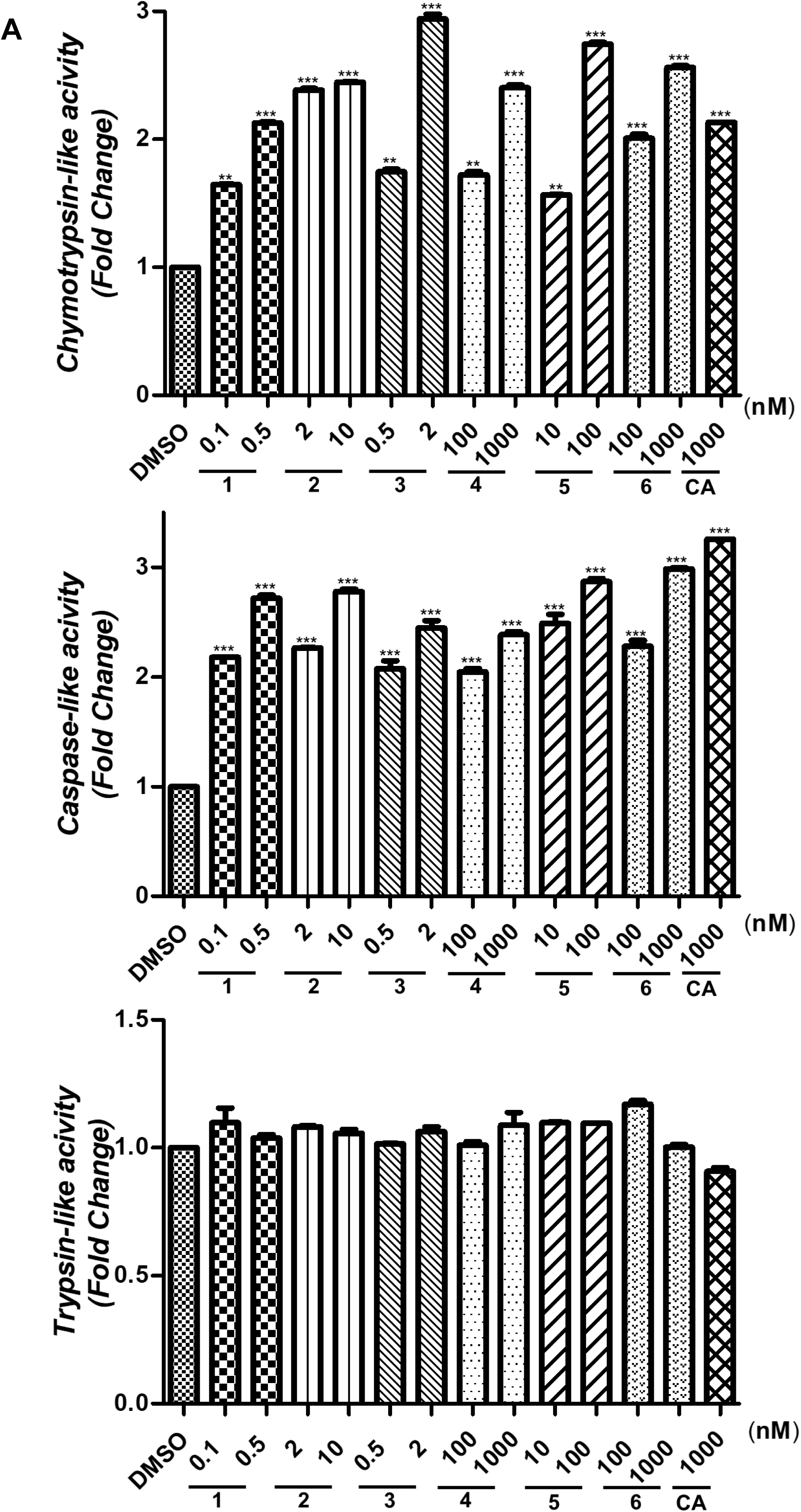
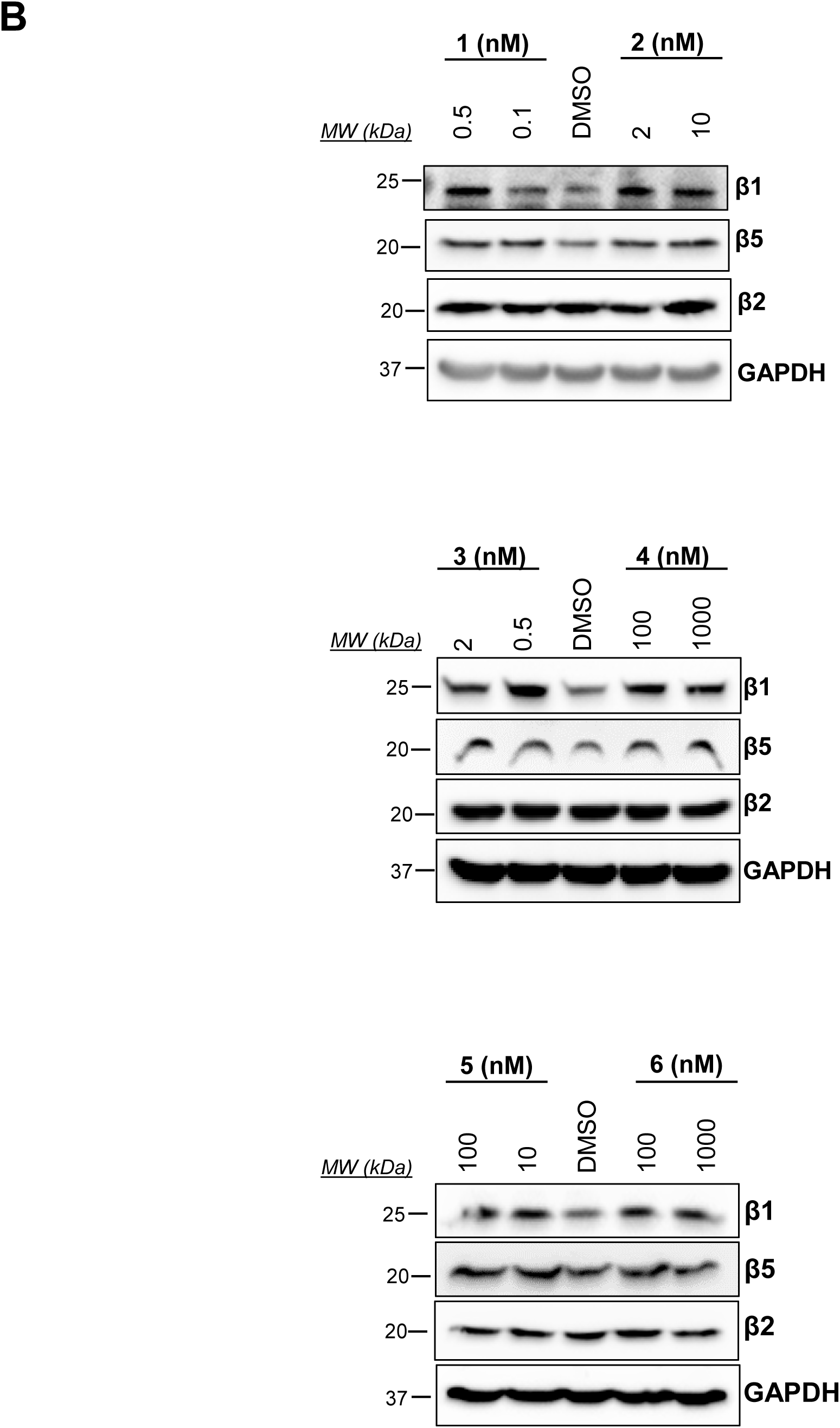
**The effect of CA derivatives on proteasome activation**. HEKn cells were treated with NRF2 active concentration of **CA** derivatives for 24 h (**A**). Fluorogenic substrates specific to each proteasome subunit were used to analyze these activities. The activity data obtained were normalized with total protein levels and graphed as fold change compared to DMSO-treated control cells. (**B**) Cellular total proteins’ levels of β1, β2, and β5 were investigated by IB. GAPDH was used as a loading control.

### 2.3. Extension of Cellular Lifespan and mitigation of replicative senescence through Novel Cycloastragenol Derivatives

The potential therapeutic properties of **CA** derivatives, such as decreasing ROS levels *via* increasing levels of cytoprotective enzymes regulated by the NRF2/ARE system, their capacity to reduce oxidative stress through increasing levels of cytoprotective enzymes via NRF2/ARE pathway and their ability to enhance proteasome activity, prompted us to investigate their effects on senescence progression. Early passage HEKn cells were continuously treated with **CA** and the derivatives at concentrations that activate NRF2. Results indicated that both **CA** (*P23, 25.71 PDL)* and its derivatives significantly extended the cellular lifespan in comparison to the control (*P19, 18.78 PDL*). Notably, metabolites **2** and **5** increased the lifespan of HEKn cells to P24 (*26.57 PDL*) and P25 (*26.81 PDL*), respectively (**Figure 4A, Supplementary Tablo S1**).

**Figure 4.**
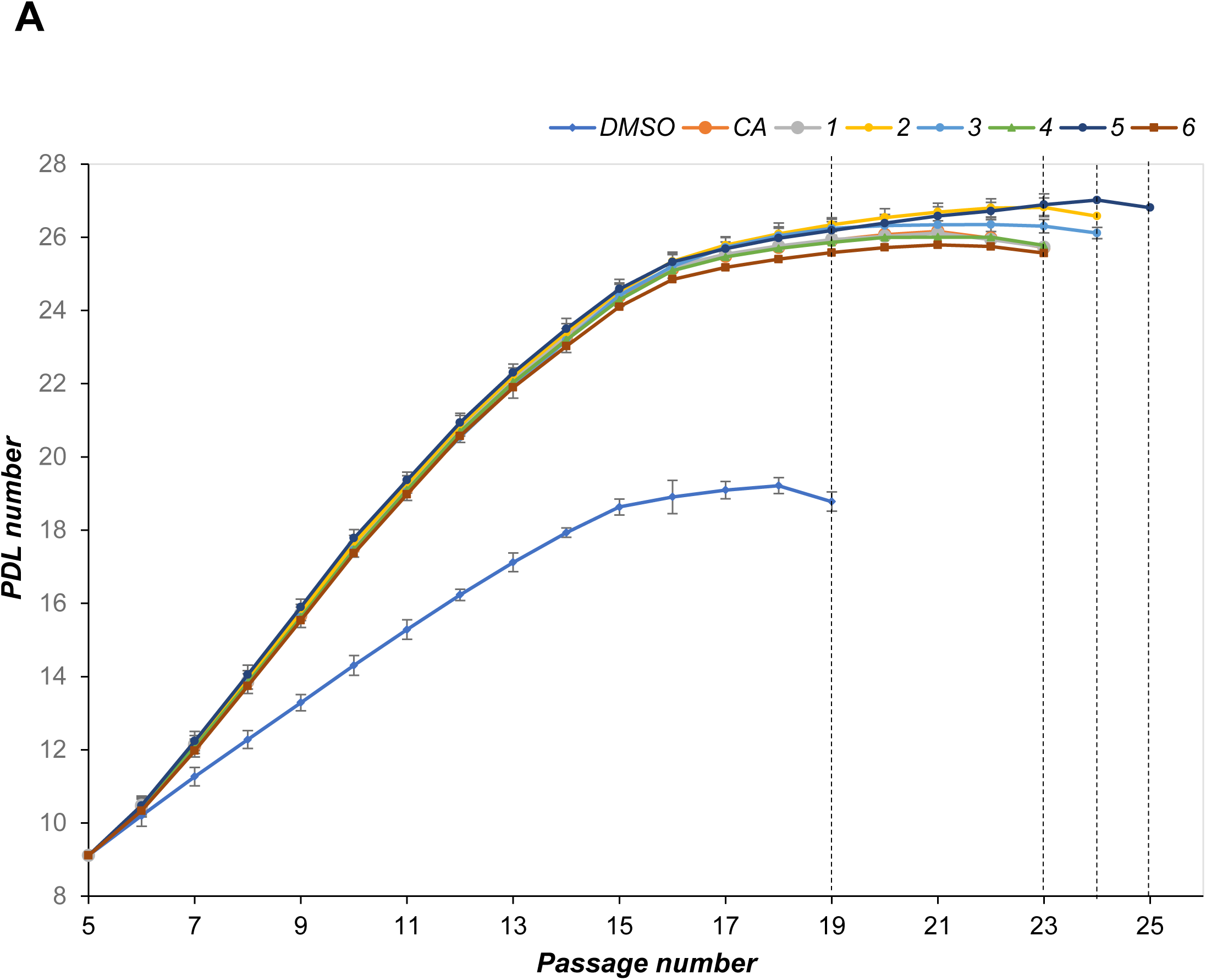
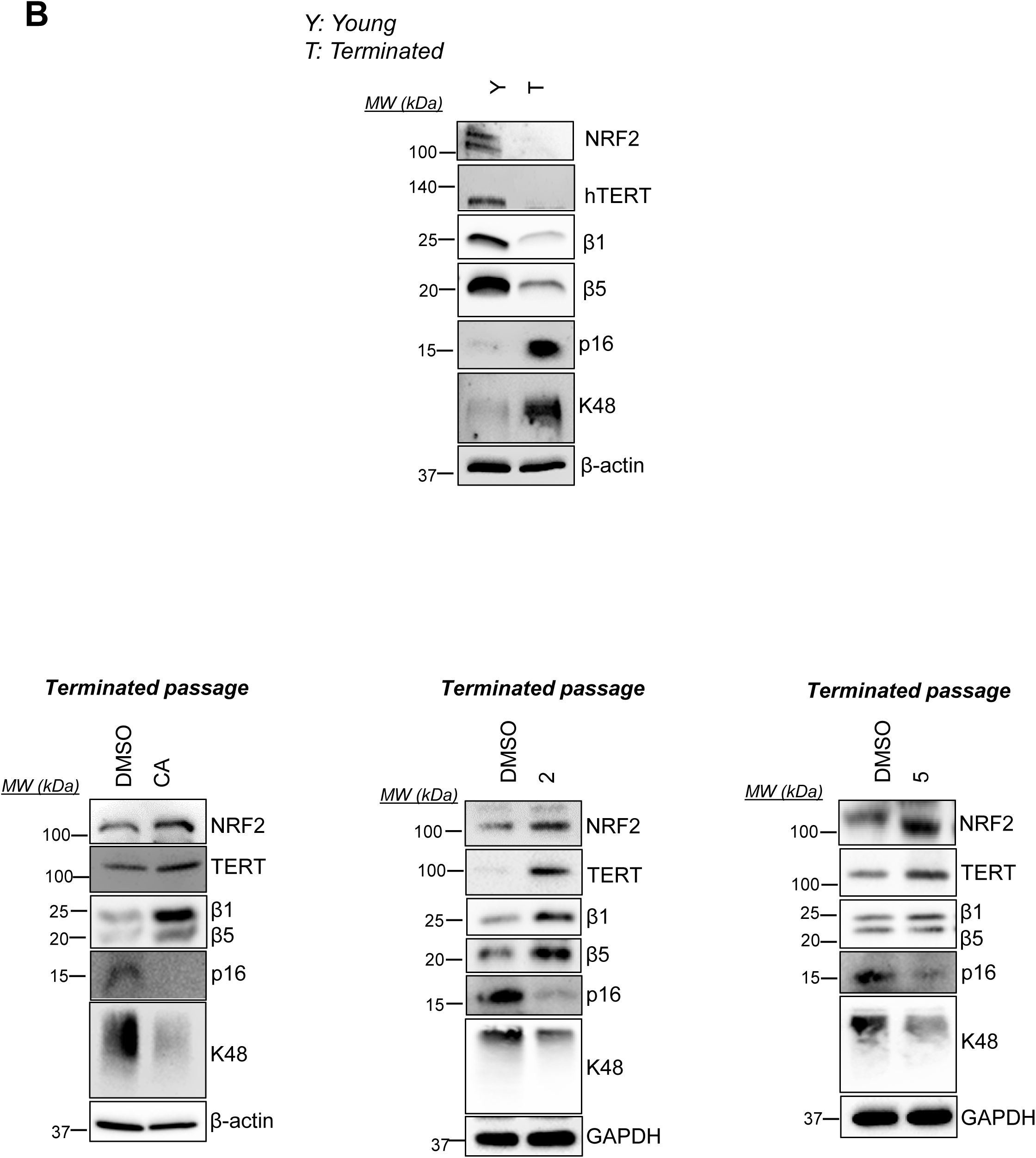
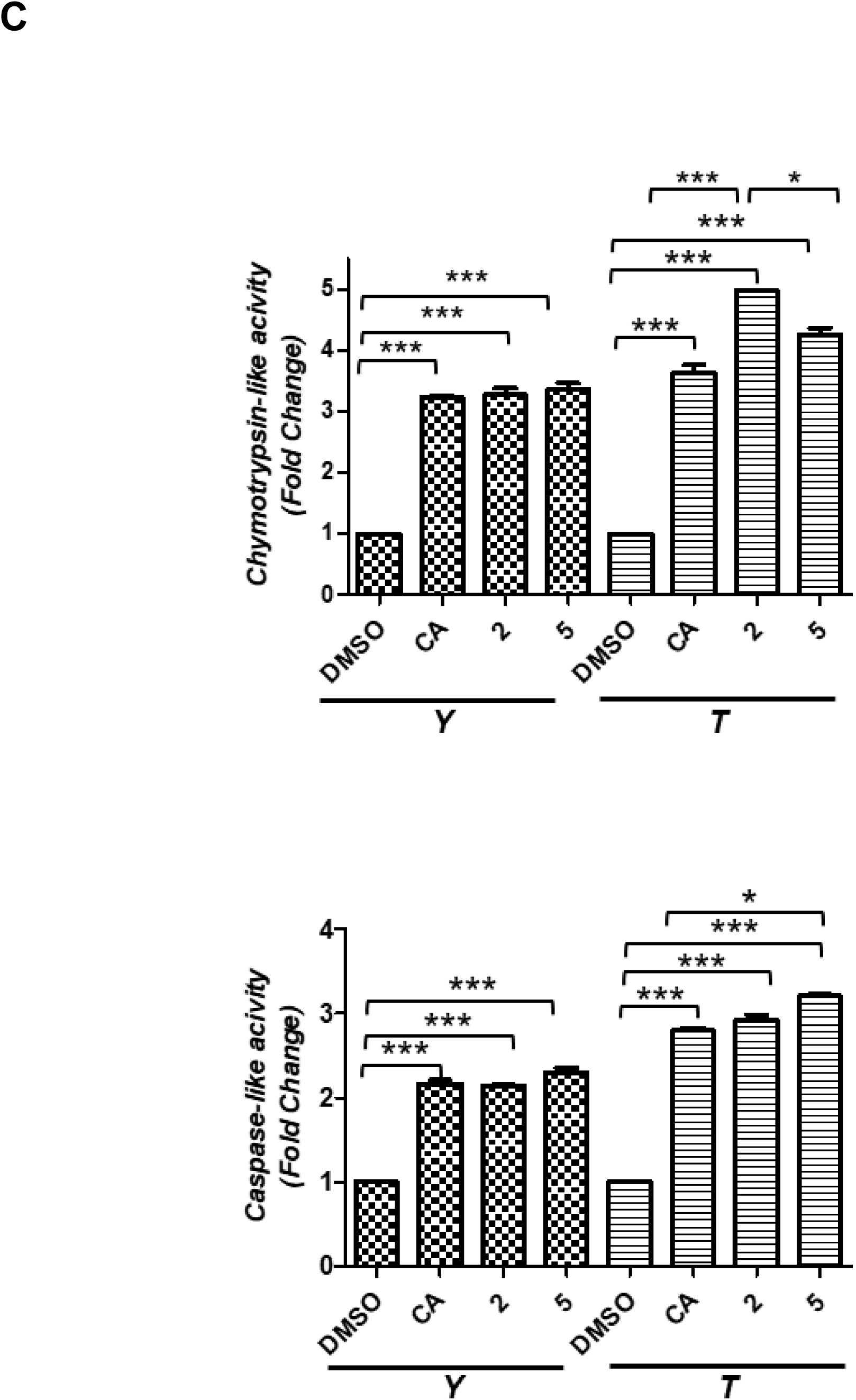
The investigation of CA and its derivatives on replicative senescence in the axis of NRF2/proteasome/telomerase systems. HEKn cells were treated with **CA** and its derivatives throughout their lifespan. (**A**) The numbers on the graph present PDL (population doubling level) at the end of the experiment. When cells reached up to 70-80% confluency, they were subcultured, and the samples taken were counted to calculate PDL. (**B**) Cellular total proteins’ levels of NRF2, hTERT, β1, β5, K48 and p16 were evaluated by IB both in young and terminated cells and terminated cells that are continuously treated with DMSO, **CA** and selected derivatives. GAPDH or β-actin was used as a loading control. (**C**) Fluorogenic substrates specific to chymotrypsin and caspase-like subunit activity were used to evaluate both young and terminated passages of HEKn cells. Total protein levels were used to normalize the obtained data. Normalized data were given as fold change compared to DMSO-treated control cells. Error bars are given as standard deviations (n = 3; *p ≤ 0.05, **p ≤ 0.001, ***p ≤ 0.005).

Given that the decline in telomere length, NRF2 and proteasomal activity is highly correlated with the replicative senescence (10, 33–37), we asked whether the stimulatory effects of **CA**, **2**, and **5** on NRF2, proteasome, and hTERT could be sustained in cells experiencing replicative aging. Starting from the first passage, cells were continuously treated with either molecules or control, and the levels of NRF2, β1, β5, and hTERT proteins were evaluated by immunoblotting, while β1 and β5 subunit activities were assessed using fluorogenic substrates in pellets taken at specific intervals. While P5 cells were used as young cells, P19 for DMSO, P23 for **CA**, and P25 for **2** and **5** were used as terminated cells. During the cellular senescence, a dramatic decline in NRF2, proteasome, and telomerase activities has been reported (34–36, 38). Consistent with this, the total protein levels of NRF2, hTERT, β1, and β5 were strikingly lower in terminated passage cells compared to young ones (**Figure 4B**). However, treatment with **CA**, as well as derivatives **2** and **5**, resulted in increased levels of NRF2, hTERT, β1, and β5 proteins in terminated HEKn cells relative to those treated with DMSO (**Figure 4C**). P16 (P16INK4A), a cell cycle regulator protein, is known to be associated with aging and senescence and is dramatically increased in senescent cells and tissues during aging or age-related diseases (39, 40). Therefore, the protein level of p16 was evaluated, and we found that its levels were very high in terminated passage cells compared to the young ones (**Figure 4B**). When cells were continuously treated with **CA** and derivatives **2** and **5** from the early passage, the levels of p16 were significantly decreased (**Figure 4B)**. During replicative senescence, an increase in poly-ubiquitinated proteins is typically observed due to the decline in proteasome activity; thus, we also evaluated K48 (K48-linkage-specific protein) levels. Low endogenous levels of K48 proteins in the young passages were contrasted by higher levels in terminated passages in DMSO-treated cells; however, K48 protein levels were observed to decrease with treatments of **CA**, **2**, and **5** in terminated passages (**Figure 4B**).

Next, we evaluated the effect of continuous treatment of CA, **2** and **5** on the proteasome β1/β5 subunit activities in terminated HEKn cells and compared it with the effect in young HEKn cells. Consistent with previous findings, CA, 2, and 5 were found to increase proteasome β1/β5 subunit activities in young passage cells (**Figure 4C)**. In terminated passage cells, while the lead compound increased β5 subunit activity by 3.64-fold, **2** and **5** significantly enhanced this activity with 4.98 and 4.26-fold increases compared to DMSO-treated cells (**Figure 4C**). Similarly, **CA** increased β1 subunit activity by 2.80-fold, and **2** and **5** had 2.93- and 3.20-fold increments compared to control cells in terminated passage cells (**Figure 4C**). Notably, continuous treatment with the compounds until the terminal passages resulted in proteasome activities that exceeded even the enhancement observed in young cells treated with the compounds. These results suggest that the delay of replicative senescence by **CA** and derivatives might be due to the axis of NRF2/proteasome/telomerase correlation.

### 2.4. Therapeutic Potential of Cycloastragenol Derivatives in Delaying Cellular Senescence through p53/p21 Modulation

Following treatment with **CA** and its derivatives, an increase in PDL of HEKn cells (up to 26,81) compared to PDL (18,71) of control cells was observed, suggesting a decrease in cellular senescence. Replicative senescence is characterized by the irreversible arrest of the cell cycle, and the p53/p21 pathway is the major pathway controlling cellular senescence, primarily activated by DNA damage and telomere attrition (41, 42). p53, cyclins, cyclin-dependent kinases (Cdk), and Cdk inhibitors (such as p21 and p27) play crucial roles in senescence signaling and are interconnected with the regulation of NRF2, proteasome and telomerase activities (43–46). When p53 is activated, the expression of p21, a cyclin-dependent kinase (CDK) inhibitor, is induced, which results in the inhibition of CDKs, essential for the transition from the G1 to the S phase of the cell cycle. Therefore, the levels of the main components of this pathway, such as p53, p21, Cyclin E, and Cdk2 proteins, were further evaluated. Consistent with the previous studies, our data revealed an increase in p53 and p21 proteins, alongside a decrease in CDK2 and Cyclin E protein levels, known to be inactivated by p21 in terminated passage HEKn cells compared to the young counterparts (**Figure 5A**). The p53 and p21 levels were shown to be significantly decreased in terminated passages following continuous treatments with **CA**, **2,** and **5** compared to DMSO-treated terminated passages. In contrast, Cyclin E and Cdk2 protein levels were markedly increased at these passages following treatments with molecules (**Figure 5B**). In summary, the findings indicate that CA and its metabolites not only counteract the biochemical hallmarks of cellular aging by upregulating the NRF2/proteasome/telomerase axis but also significantly modulate key senescence-associated signaling pathway, p53/p21 pathway, suggesting their potential as therapeutic agents in the mitigation of replicative senescence.

**Figure 5.**
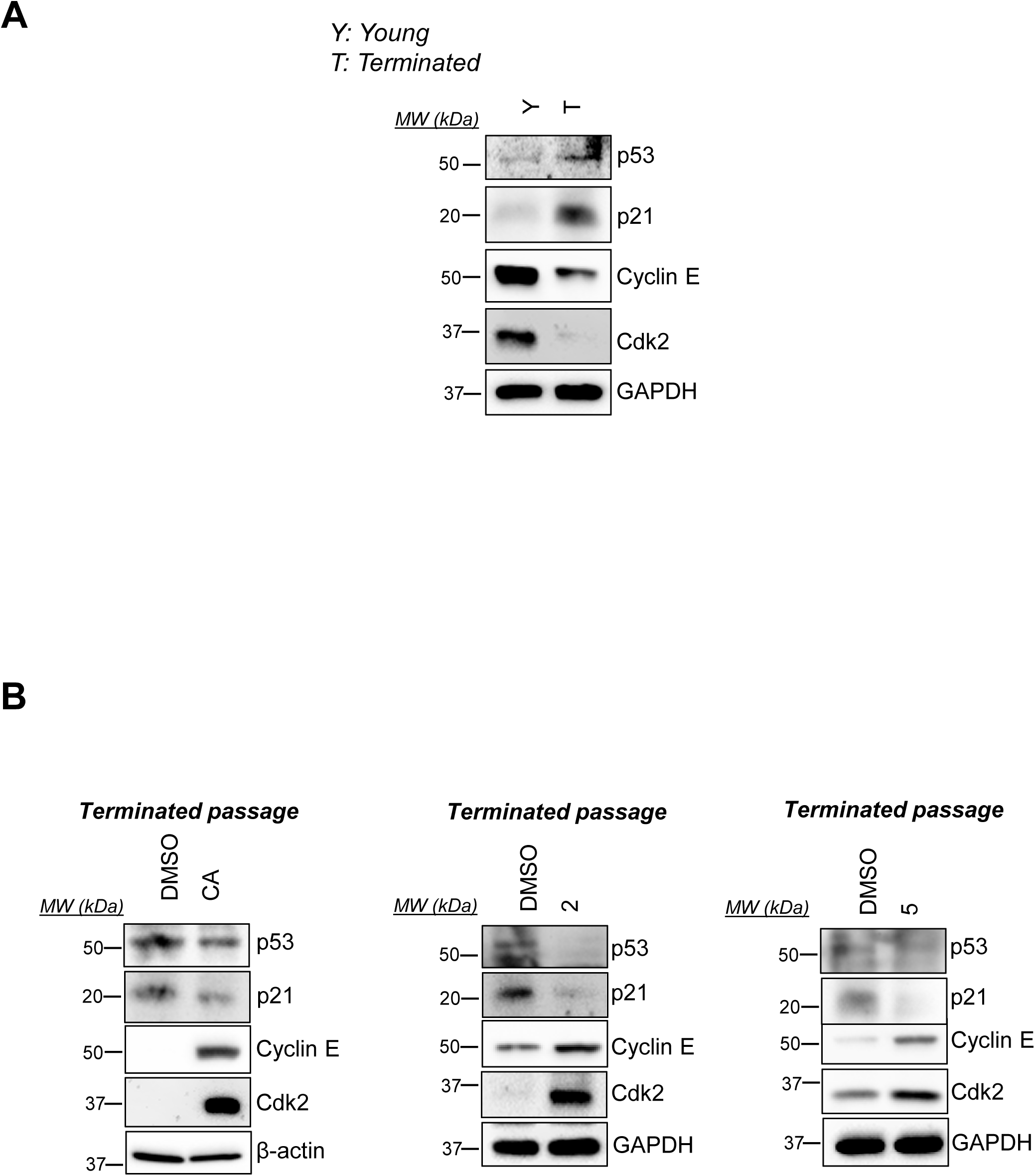
The role of Cycloastragenol Derivatives in delaying Cellular Senescence via p53/p21 pathway. HEKn cells were treated with **CA** and its derivatives throughout their lifespan. (**A**) Cellular total proteins’ levels of p53, p21, Cyclin E and Cdk2 were evaluated by IB both in DMSO-treated young and terminated cells. GAPDH was used as a loading control. (B) Cellular total proteins’ levels of p53, p21, Cyclin E and Cdk2 were investigated by IB in DMSO-treated and compounds treated terminated cells. GAPDH or β-actin was used as a loading control.

### 2.5. The Role of CA Derivatives in Preventing Glutamate-Induced Neurotoxicity

Glutamate, the central nervous system’s primary excitatory neurotransmitter, is essential for regulating post-synaptic glutamate receptor activation during normal physiological neurotransmission (47). Elevated levels of glutamate can lead to excessive accumulation of NMDA (N-methyl-D-aspartate) type glutamate receptors, resulting in neuronal cell death through a consequence of a serious of cascade events such as membrane depolymerization, excessive Ca+2 ion influx, protease activation, and oxidative stress (47–50). It has been observed that particular cell signaling pathways may play a role in both glutamate excitotoxicity and replicative senescence. The p53/p21 pathway, which plays a crucial role in cell cycle regulation, is one such pathway that can be activated in response to DNA damage under conditions of both glutamate excitotoxicity and senescence. Therefore, in our current study, we have assessed the potential protective effects of **CA** and its biotransformation products on glutamate-induced neurotoxicity in HCN-2, followed by the initial study that determined the optimal glutamate concentration (**Supplementary S4**).

While cell viability decreased to 50.04% in control cells exposed to glutamate, pre-treating cells with **CA** at 10-100 and 1000 nM concentrations for one hour prior to glutamate exposure improved cell survival rates up to 60.9%, 67.8%, and 73.5%, respectively (**Figure 6A**). Remarkably, all **CA** derivatives also protected cells against glutamate toxicity, with some metabolites showing protection at even lower concentrations than **CA** itself. A similar protective effect of CA and its derivatives was also observed in glutamate-induced neurotoxicity in primary rat cortex neurons (**Supplementary S5**).

**Figure 6.**
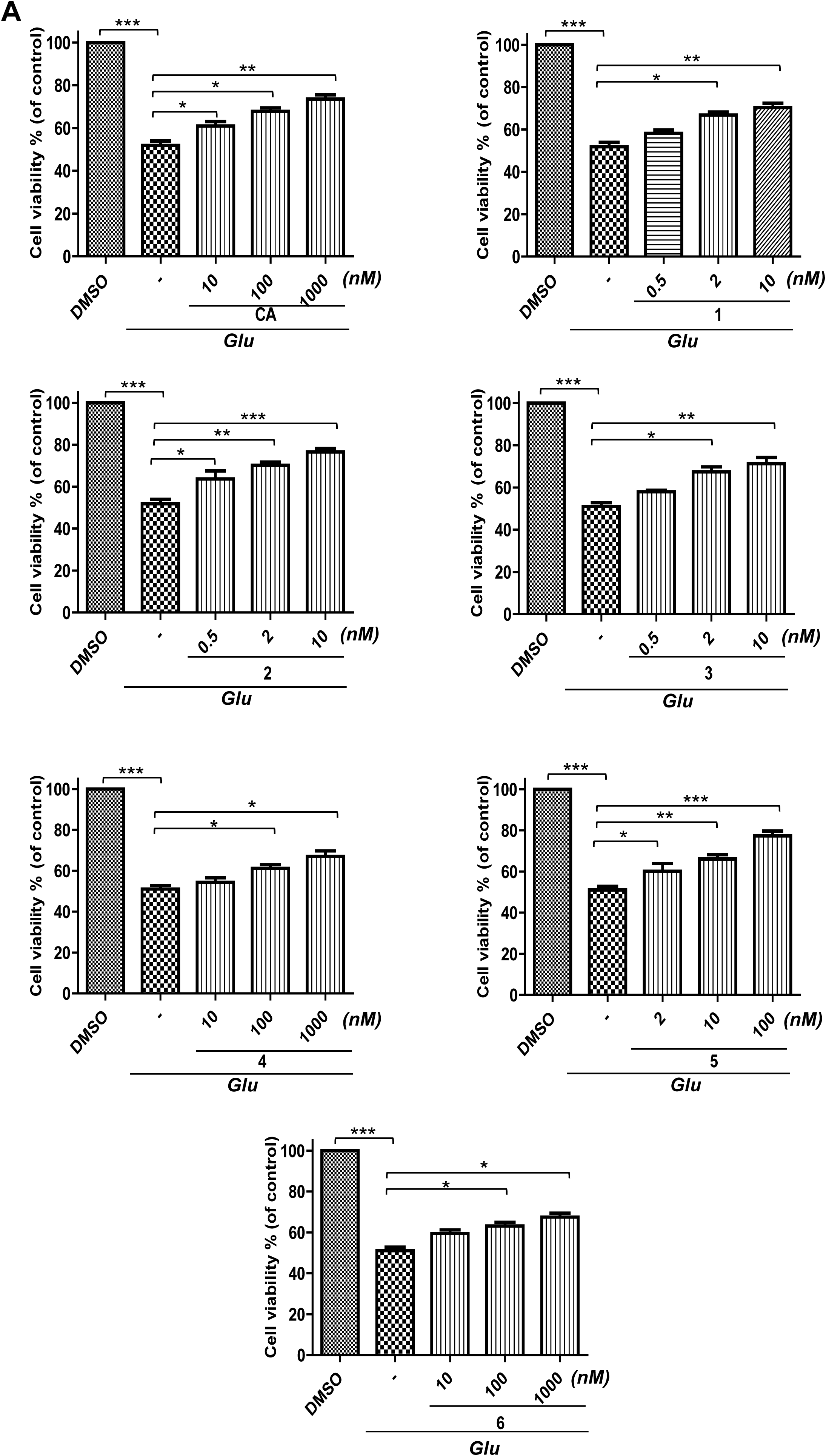
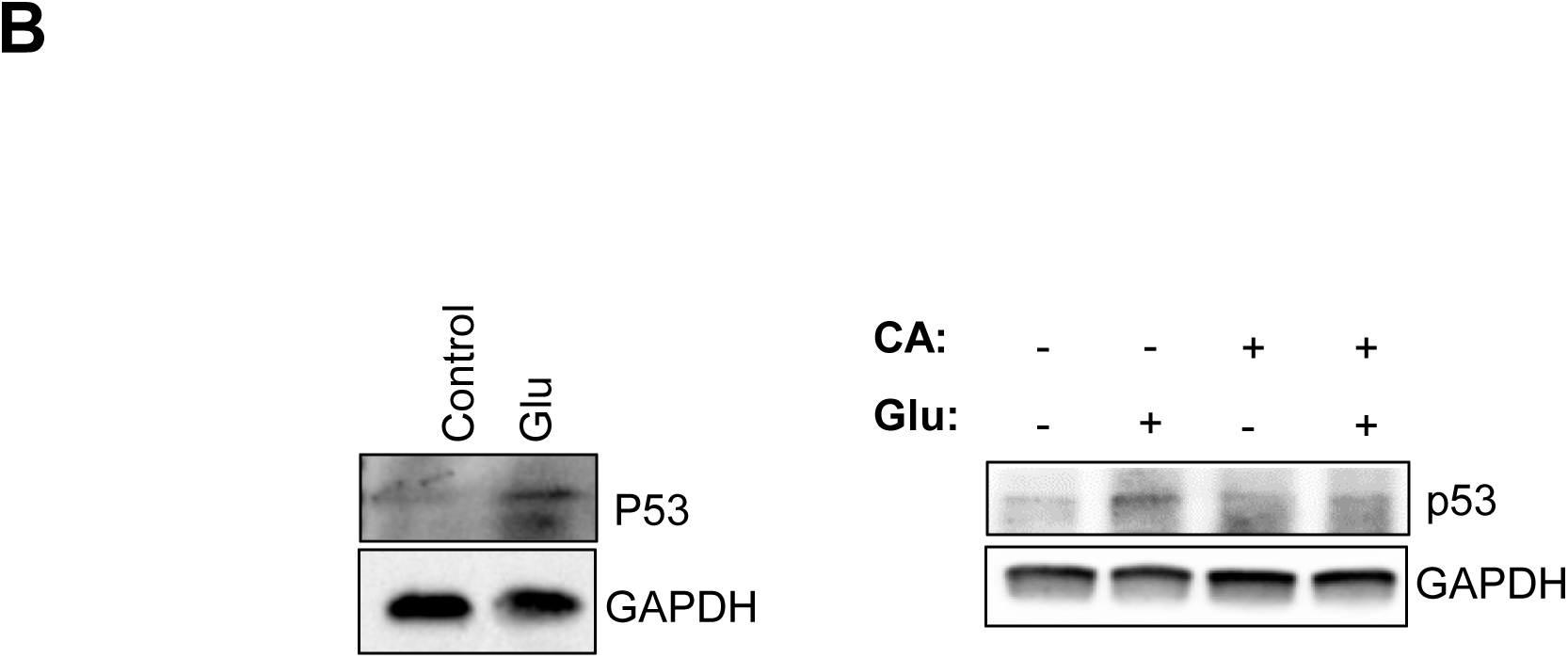
The neuroprotection of CA and its derivatives against glutamate-induced neurotoxicity in HCN-2 cells. HCN-2 cells were pre-treated with **CA** and its derivatives for 1 h, then treated with 500 μM glutamate for 24 h**. (A)** The viability of cells was analyzed with an MTT reagent. The absorbance value of DMSO-treated cells was considered as 100%, and then the viabilities of **CA** and its derivatives were calculated according to DMSO. The experiment was conducted in triplicate. To determine the significance of the differences, a Student’s t-test was used. Error bars are showed as standard deviations (n = 3; *p ≤ 0.05, **p ≤ 0.001, ***p ≤ 0.005). (**B**) Total protein levels of p53 were determined by IB. GAPDH was used as a loading control.

A series of papers have contributed to unraveling the contribution of p53 to neurodegeneration, and among several types of neuronal toxicity, glutamate excitotoxicity was also reported to upregulate the transcriptional activity of p53 to induce pro-apoptotic pathway (51, 52). Consequently, the downregulation of p53 protects neurons from cellular death (53). As we observed the protective effect of CA and derivatives against glutamate-induced neurotoxicity, the effects of **CA** on p53 protein levels were evaluated in this toxicity model. Results revealed that the p53 protein level increased with glutamate treatment and decreased with **CA** pre-treatment (**Figure 6B)**.

## 3. Discussion

In traditional Chinese medicine, various herbal products have been used for anti-aging therapies for centuries. One of the most significant herbs is *Astragalus membranaceus*, known for its long medical history and mention the complete Pharmacopoeia of China. Initial findings demonstrated the potency of the extract of this plant in delaying replicative senescence in human diploid fibroblast, leading to efforts to identify the compounds responsible for this activity. Among the significant metabolites isolated from *A*. *membranaceus* was Astragaloside IV (AST IV), which has anti-aging properties that protect skin cells against UVA irradiation *via* suppressing MAPK and NK-κB pathways and regulating the TGF-β/Smad pathway (54, 55). Also, AST IV exhibits immunomodulatory effects without toxicity, increasing B and T cell proliferation and dramatically reducing IL-1 and TNF-α levels (56, 57).

Additionally, AST IV possesses protective properties against ROS (58, 59), cardiovascular diseases (60), myocardial (61), or cerebral ischemia-reperfusion injury (62, 63) via enhanced NRF2 activation. Another critical compound derived from AST IV is Cycloastragenol (**CA**), which also exhibits potent anti-aging properties. Several studies indicate that **CA** affects several cellular processes, including enhancement of wound healing, inhibition of autophagy in airway inflammation models (64), induction of CREB activation in neuronal cells, and exertion of anti-depressant effect by enhancing neurotrophic signaling (65). **CA** also upregulates SIRT1, preventing neuroinflammation in the ischemic brain (20), and is hepatoprotective in liver fibrosis induced by carbon-tetrachloride (17).

Moreover, **CA** augments telomere length and life span in mice without increasing cancer incidence (66). Strikingly, **CA** demonstrates anti-cancer properties by activating p53 in colon cancer cell lines (67) and downregulating cathepsin B, thus enhancing anti-tumor immunity through improved MHC-I presentation (68). TA-65®, a commercial formulation of **CA**, has been included in clinical trials to evaluate its effects on several conditions, such as Alzheimer’s disease and metabolic syndrome (69, 70).

In this study, we evaluated the effect of these telomerase-active derivatives (**Figure 1**) on the NRF2/proteasome axis and replicative senescence in human epidermal keratinocytes (HEKn). Our findings revealed that biotransformation products increased NRF2 expression levels, nuclear translocation, and cytoprotective enzyme levels, such as HO-1, GCLC, and GR, attributed to elevated ARE promoter activity (**Supplementary S2B-E, Figure 2)**. This conferred protection against oxidative stress induced by H_2_O_2_ (**Supplementary S2F)**. When treated with increasing concentrations of the biotransformation products, these novel derivatives showed NRF2 activation at varying concentrations. Derivative **1**, possessing the α- oriented hydroxyl group at C-12, enhanced nuclear NRF2 levels at lower concentrations (0.1 and 0.5 nM) with 2.97 and 3.47-fold change compared to DMSO-treated cells as a control (**Figure 2A)** (71) (26). Derivative **2**, a product of the Baeyer-Villiger monooxygenase enzyme and containing a 7-membered lactone in the A ring, enhanced the protein level at 2 and 10 nM with 2.61- and 2.50-fold, respectively (**Figure 2A**). When examining the 3-oxo derivatives, derivative **3** increased the nuclear transcriptional activity of NRF2 at 0.5 and 2 nM concentrations with 2.76- and 3.25-fold, whereas **6** increased this activity at 100 and 1000 nM concentrations with 1.37- and 1.69-fold (**Figure 2A**). When 3(4)-seco derivatives (**4** and **5**) were tested, **4** augmented this at 100 and 1000 nM with 2.14- and 2.50-fold, but **5** provided 1.64- and 2.02-fold change at 10 and 100 nm, respectively (**Figure 2A**). Our previous report indicated that **CA** increased NRF2 expression and nuclear transcriptional activity, leading to enhanced protection against oxidative stress at higher concentrations such as 100- and 1000 nM (25). This study demonstrates for the first time that these novel derivatives are potent NRF2 activators, increasing cytoprotective enzyme levels by enhancing ARE-promoter activity (**Figure 2C, Supplementary S2E**). Consistent with findings in young passage HEKn cells, all derivatives increased nuclear NRF2 transcriptional activity, resulting in elevated levels of GR, HO-1, and GCLC enzymes, protecting senescent cells from oxidative stress (**Supplementary S2A and S2E**). Notably, derivatives **1**, **2**, **3**, and **5** showed higher potency at lower concentrations, suggesting improved bioavailability and target interaction compared to **CA**. Therefore, monooxygenation at the C-12 position, oxidation, lactone formation, ring opening, and dehydration reactions in the A-ring of **CA** or **AG** might be essential for NRF2 activation, resulting in transcriptional regulation of both cytoprotective enzyme and proteasome subunit genes. However, further screening with more biotransformation products with similar modifications is required to elucidate the complete structure-activity relationship thoroughly.

One of the downstream targets of the NRF2/ARE system is proteasome subunit genes and activities (72–74). Furthermore, the telomerase enzyme subunit TERT has a novel non-canonical role unrelated to telomere elongation as a chaperone protein in promoting proteasome assembly/activation (75). Targeting the proteasome with small compounds has gained momentum for treating or preventing age-related diseases and promoting healthy aging. Several natural compounds were reported as proteasome activators, such as Hederagenin, Sulforaphane (76), Betulinic acid (77), Oleuropein (78), 18α-Glycyrrhetinic acid (34), and Ursolic acid (79). We also previously reported that **CA** is a novel proteasome activator, with its activation induced by telomerase activity mediated by NRF2 (25). Building on these findings, we evaluated the effects of CA derivatives on proteasome status in the present study. We identified that these derivatives increase both β1 (responsible for caspase-like activity) and β5 (responsible for chymotrypsin-like activity) subunit activities and protein levels but do not induce a significant change in the β2 subunit (trypsin-like activity) (**Figure 3A** and **B**). Additionally, continuous treatment with the compounds until the terminal passages resulted in proteasome activities that exceeded even the enhancement observed in young cells treated with the compounds (**Figure 4C**).

Replicative senescence, a central hallmark of aging, involves several biological processes, including chromatin remodeling, metabolic reprogramming, and the release of SASP (senescence-associated secretory phenotype) factors (80–83) and is characterized by a permanent arrest of cell proliferation (84). This process is heavily influenced by telomere shortening, oxidative stress, and the decline in proteostasis (34, 38, 85, 86). These factors collectively contribute to cellular aging by triggering pathways that involve key regulators such as p53 and p21. For instance, telomere shortening triggers a DNA damage response (DDR) involving ATM, p53, and inducing cell cycle inhibitors p21 and p16, which arrest cell proliferation and maintain cellular senescence. Telomere dysfunction-activated DDR, rather than telomere dysfunction itself, is reported to cause cellular senescence, which facilitates the age-related loss of tissue functions through the secretion of SASP factors (4, 5). Our findings indicate that **CA** derivatives can counteract this process not only by enhancing telomerase activity but also by reducing the activation of p53 and its downstream effector p21 (**Figure 5B)**. This reduction in p53 and p21 levels alleviates the cell cycle blockade, evidenced by the prolonged lifespan of HEKn cells treated with **CA** derivatives compared to control cells (**Figure 5B**). The interplay between p53 and NRF2 further elucidates the mechanism behind the protective effects of **CA** derivatives. NRF2 has been shown to interact with p53, where NRF2 activation can suppress p53-mediated cellular senescence pathways (87, 88). It has also been shown that the inhibition of p53 can enhance NRF2 signaling, thereby promoting antioxidant defenses and reducing cellular senescence.

This interaction highlights a potential mechanism through which CA derivatives can exert their anti-senescence effects, as the **CA** derivatives likely modulate the p53/NRF2 axis, enhancing NRF2 activity and providing robust protection against oxidative stress. Interestingly, the partial inhibition of proteasomes has been shown to trigger premature senescence through the p53 pathway (13), and the role of p53 in regulating proteasome inhibition-mediated senescence adds another layer of complexity to our findings. Studies have shown that functional p53 is essential for maintaining senescence upon proteasome inhibition, whereas cells lacking p53 can escape senescence and resume growth (13). Our derivatives’ ability to modulate p53, along with NRF2 and proteasome activities, supports the hypothesis that these molecules can delay replicative senescence by stabilizing key regulatory pathways. Furthermore, our results indicated that **CA** and its derivatives protect against glutamate-induced excitotoxicity, which is known to upregulate p53 and lead to neuronal cell death (**Figure 6A**). **CA** reduced p53 protein levels in glutamate-treated cells, providing neuroprotection (**Figure 6B**). This suggests that **CA**’s modulation of the p53 pathway contributes not only to delaying replicative senescence but also to protecting against neuronal damage. The combined action on p53, NRF2, and proteasome pathways underscores the potential of **CA** and its derivatives in both anti-aging and neuroprotective therapies.

In summary, the novel **CA** derivatives demonstrate multifaceted benefits by enhancing NRF2 activity, maintaining proteostasis, and modulating the p53 pathway. These combined actions contribute to the delay of replicative senescence and improved cellular longevity. The observed enhancement of telomerase activity and subsequent delay in senescence further support the potential of **CA** derivatives as multi-targeted agents for increasing lifespan with healty aging and preventing age-related diseases. Continued research on these derivatives may yield significant insights into the development of comprehensive anti-aging interventions.

## 4. Material and Methods

### 4.1. Reagent and Antibodies

In this study, the proteasome fluorogenic substrates Suc-LLVY-AMC-BML-P802-0005 for the ChT-L (chymotrypsin-like), Z-LLE-AMC-BML-ZW9345-0005 for the C-L (caspase-like), and Boc-LRR-AMC-BML-BW8515 for the T-L (trypsin-like were purchased from Enzo Life Sciences. While primary rabbit antibodies as NRF2 (Abcam, Ab-62352), NRF2 (Bioss, bsm- 52179R), HO-1 (CST-5853), c-JUN (CST-9165), p16 (CST, 80772), K48 (CST, 8081) and β5 (BML-PW8895) were used, primary mouse antibodies as β1 (BML-PW8140), β2 (BML- PW8145), Cdk-2 (sc-6248), p27 (sc-1641), p21 (sc-6246), Cyclin E (sc-377100), p53 (CST, 2524) were used in this study. Antibodies against β-actin (Sigma-Aldrich-A5316), GAPDH (CST, 5174, and Lamin A-C (CST, 4777) were used as loading controls. Secondary antibodies (Goat anti-rabbit-31460 and Goat anti-mouse-31430) were purchased from Thermo Fisher Scientific. DMSO (Dimethylsulfoxide; D2650) was purchased from Sigma Aldrich.

The derivatives were obtained via biotransformation from the previous studies and used in the present study (26–28).

### 4.2. Cell lines & Culture Conditions

HEKn (human neonatal epithelial keratinocyte cells) obtained from American Type Culture Collection (ATCC; PCS-200-010) were maintained and cultured in Dermal Cell Basal Media (ATCC; PCS-200-030) including Keratinocyte Growth Kit (ATCC; PCS-200-040) according to the manufacturer’s instructions. When the confluency of HEKn cells reached %70- 80, they were passaged and seeded at a density between 2500 to 5000 cells per cm2.

Human neuron cortical cells (HCN-2, ATCC, CRL-3592) were cultured in DMEM (Lonza, BE12-604F) supplemented with 4 mM L-glutamine (Biological Ind., 03-020-1B) and 10% FBS (Biological Industries, 04-007-1A). When these cells reached 80% confluency, cells were subcultured at a 1:3 ratio.

Primary rat cortex cells (Gibco, A10840-02) were cultured in Poly-D-lysine coated 96 well and 12 well plates. In brief, the cells were maintained in Neurobasal™ Plus Medium (Gibco, A3582901) supplemented with B-27™ Plus Supplement (Gibco, A3582801) and Glutamine (Sigma, G7513). Half of the media was refreshed every third day with fresh media.

### 4.3. Cellular fractionation and NRF2 Transcription Factor assay

HEKn cells were treated with increased concentrations of **CA** derivatives for 24 h. Nuclear-cytoplasmic fractionations were conducted according to the manufacturer’s guidelines (NE-PER, Pierce, Thermo Fisher Scientific, US, 78833). Then, nuclear lysates were assessed to ELISA for NRF2 Transcription Factor assay (Cayman Chemical, US, 600590) to investigate whether these derivatives affect the transcriptional activity of nuclear NRF2. Cytoplasmic and nuclear lysates were also used for nuclear localization for NRF2 *via* immunoblotting (IB). In IB experiments, while GAPDH was used as a cytoplasmic loading control, Lamin A-C was used as a nuclear loading control.

### 4.4. Luciferase activity assay

HEKn cells were transfected with pGL4.37[luc2P/ARE/Hygro] Vector (Promega, US, E364A) to evaluate whether derivatives affect ARE promotor activity for 24 h. Then, HEKn cells were treated with NRF2 active concentrations of derivates for 24 h. Next, cells were lysed in Cell Culture Lysis Reagent (Promega, US, E1500). The lysates were used to determine luciferase activity assay according to the manufacturer’s instructions. Data were presented as fold-change compared DMSO-treated cells with standard deviations (SD) and means (n:3).

### 4.5. Measurement of Cellular ROS levels

2’-7’-Dichlorodihydrofluorescein diacetate (H_2_DCFDA) (Enzo; ALX-610-022-M050) was used for the determination of cellular ROS level as previously described (89). For this purpose, HEKn cells were pre-treated with NRF2 active concentrations of derivatives for 1h. After 1 h pre-treatments of them, cells were treated with 250 µM H_2_O_2_ for 24 h. H_2_DCFDA, suspended in 1X PBS to reach 10 µM concentration, was added to cells and incubated at 37

°C for 30 min (34). After the incubation period, half of the cells were used to measure ROS levels at 495 and 520 nm using a microplate reader (Varioskan, Thermo Fisher Scientific). The other half was used to determine protein concentrations to normalize data. This experiment was done in three independent experiments.

### 4.6. Replicative senescence

HEKn cells were treated with CA and derivatives every other day and passaged when the cells reached 70-80% confluency. The day after passaging, the medium was renewed to prevent trypsin’s negative effect, and the molecules were re-treated the following day.

The HEKn with the youngest passage number in our cell bank is passage 5; studies started with passage 5.

During every passage, cells were counted, and PDLs (*population doubling levels*) were calculated as in the formula below (90):

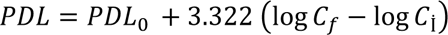

*PLD_0_: initial population doubling level*

*C_İ_: initial cell number seeded into a culture plate*

*C_f_: final cell number*

The methods used for further analysis are described in detail in Sections 2.9 and 2.10.

### 4.7. Proteasome activity assays

The proteasome subunits’ activities were determined as previously described (91). Briefly, HEKn cells were treated with NRF2 active concentrations of derivatives for 24 h (91). The cells were lysed in the buffer containing 8,56 g sucrose, 0.6 g HEPES, 0.2 g MgCl_2_, 0.037 g EDTA and freshly prepared 1 mM DTT. After incubating lysates with reaction mixture buffer (involving .5 mM MgOAc, 7.5 mM, MgCl_2_, 45 mM KCl, and 1 mM DTT), fluorogenic substrates (specific to each subunit) were incubated for 1 h. The measurements were done by a fluorescent reader at 360 nm excitation/460 nm emission (Varioscan, Thermo Fisher Scientific, US). DMSO was used as an experimental control. Normalization was performed by total protein concentrations determined via BCA (bicinchoninic acid) assay (Thermo Fisher Scientific, USA). Data were given as fold-change compared DMSO-treated cells with SDs and means (n:3).

### 4.8. Immunoblotting (IB)

RIPA buffer (1% Nonidet P-40, 0.5% sodium deoxycholate, and 0.1% SDS in 1X PBS, pH 8.0 with protease inhibitor cocktail) was used for obtaining cell lysates. Protein concentrations were determined by BCA assay. Equal amounts of samples were prepared and denatured in 4X Laemmli buffer at 95 degrees. After denaturation, samples were loaded into SDS-PAGE gels and transferred to PVDF membranes (EMD Millipore, US, IPVH00010). After blocking in 5% non-fat dry milk prepared in 1X PBS-0.1% Tween-20, membranes were incubated first as primary antibodies (listed in Material section), then secondary antibodies (listed in Material section). While Lamin A-C was used as a nuclear loading control, GAPDH and β-actin were used as a cytoplasmic or whole lysate loading control. Chemiluminescence signals were identified with Clarity ECL substrate solution (BIORAD, US, 1705061) by Vilber Loumart FX-7 (Thermo Fisher Scientific, US). Western blot images were analyzed with ImageJ software (http://imagej.nih.gov/ij). IB experiments were performed at least in three independent replicates.

### 4.9. Cell viability assay

To determine the IC_50_ value of Glutamate, cells were treated with increased concentrations (0-1000 µM) for 24 h. Then, the mixture of MTT and medium (1:9) was replaced with old media. The formazan crystals were dissolved in DMSO. The absorbance was measured with a microplate reader at 570 nm (Varioscan, Thermo Fisher Scientific, US) and 690 nm as a reference wavelength. Cell viability was presented as a percentage of cell viability compared to DMSO-treated cells as control cells. The experiment was conducted with three different replicates. After determining the IC_50_ value of Glutamate, to evaluate whether **CA** and its derivatives have any protection against glutamate-induced excitotoxicity or not, HCN-2 and rat cortex cells were pre-treated with them for 1 h and then treated with Glutamate (500 µM) for 24 h. after incubation period completed, MTT assay was performed described before. Data were given as fold-change compared DMSO-treated cells with SDs and means (n:3).

### 4.10. Statistical Analysis

Data obtained were presented as means± standard deviation (SD). Student t-test or One-way ANOVA Post Hoc test was used for statistical analysis using GraphPad Prism software. The significance of the differences was given as *p ≤ 0.05, **p ≤ 0.001, ***p ≤ 0.00

## Author Contributions

P.B.K and E.B. conceived the hypotheses and designed this study. S.Y. participated in the design of the experiments, conducted the experiments, and collected the data. All authors analyzed the results, contributed to the manuscript preparation and read/approved the manuscript.

## Data statement availability

The data supporting this study’s findings are available from the corresponding author upon request.

## Acknowledgment

We thank the Pharmaceutical Sciences Research Centre (FABAL, Ege University, Faculty of Pharmacy) for equipment support and Bionorm Natural Products for donating **CA**. We would like to thank Dr. Melis Küçüksolak for her support to isolate and characterize **CA** derivatives.

## SUPPLEMENTARY MATERIALS

**Figure S1. CA and its derivatives increased hTERT protein levels in HEKn cells.** HEKn cells were treated with NRF2 active concentrations of CA and derivatives for 24 h. The protein levels of hTERT were analyzed by IB. GAPDH was used as a loading control.

**Figure S2. The effects of CA derivatives on NRF2/ARE systems.** (**A**) **CA** derivatives enhanced nuclear NRF2 transcriptional activity in senescent HEKn cells. HEKn cells were treated with the indicated concentration of derivatives for 24 h. After the fractionation, the nuclear lysates were used to determine nuclear NRF2 transcriptional activity with ELISA. The data were normalized with nuclear proteins level, then presented as fold change compared to the DMSO-treated control cells. Error bars are presented as standard deviations (n = 3; *p ≤ 0.05, **p ≤ 0.001, ***p ≤ 0.005). (**B**) Cellular localization of NRF2 was also investigated by immunofluorescence analysis using an anti-NRF2 antibody. DAPI was used for nuclei staining. (C) NRF2 mRNA levels were evaluated by RT-qPCR. Error bars are presented as standard error (n=3, *p ≤ 0.05, **p ≤ 0.001, ***p ≤ 0.005). (**D**) NRF2 and c-JUN protein levels were investigated by IB. GAPDH was used as a loading control. (**E**) HO-1, GCLC, and GR cytoprotective enzyme levels were analyzed by ELISA, both in young and old HEKn cells. Each enzyme levels were normalized to total protein levels and given as a fold change compared to DMSO-treated cells. Error bars are given as standard deviations (n = 3; *p ≤ 0.05, **p ≤ 0.001, ***p ≤ 0.005).

**Figure S3. CA derivatives enhanced the proteasome activation in senescent HEKn cells.** Senescent HEKn cells were treated with NRF2 active concentrations of derivatives for 24 h. Substrates specific to caspase and chymotrypsin-like subunit activity were used to determine these activities. The obtained data were normalized with total protein levels and presented as fold change compared to the DMSO-treated control cell.

**Figure S4. Identification of the optimum concentration of glutamate concentration in HCN-2 cells.** HCN-2 cells were with increased concentrations of glutamate for 24 h. MTT reagent was used to determine cell viability. DMSO-treated cells were considered 100%, and the others were calculated according to DMSO-treated cells. The experiment was performed in triplicate. Student’s t test was used for determination of significance of the differences. Error bars are showed as standard deviations (n = 3; *p ≤ 0.05, **p ≤ 0.001, ***p ≤ 0.005).

**Figure S5. CA and its derivatives protect rat cortex cells against glutamate excitotoxicity.** Rat cortex cells were pre-treated with CA and its derivatives for 1h and then treated with glutamate (500 μM) for 24 h. MTT reagent was used for cell viability. DMSO-treated cells were considered as 100% cell viability. This experiment was conducted in triplicate. To determine the significance of the differences, a Student’s t-test was used. Error bars are given as standard deviations (n = 3; *p ≤ 0.05, **p ≤ 0.001, ***p ≤ 0.005).

Tablo S1. The effect of CA and its derivatives on PDLs (Population doubling levels).

Tablo S2. q-RT-PCR primer list.

## Methods

### Immunofluorescence (IF) Analysis

HEKn cells were seeded on 6 well plates with a coverslip to evaluate nuclear NRF2 localization from cytoplasm *via* fluorescence microscopy (Olympus IX70, Japan) (92). The next day, cells were treated with NRF2 active concentrations of derivatives for 24 h. According to our previous study, IF steps were performed (25). Briefly, cells were fixed with 4% paraformaldehyde in PBS for 30 min at 4°C. Then, cells were washed six times with 1X PBS. After permeabilization and blocking were completed, cells were incubated first with antibody against NRF2 (Abcam, Ab-62352, 1:250 dilution) at 4°C overnight and next incubated with seconder antibody suitable for IF (Pierce, A-11008, 1:500 dilution). Mounting was conducted with DAPI (Invitrogen, P36965) for nuclei staining.

### Cytoprotective enzyme activity assay

To determine cytoprotective enzyme levels such as HO-1, GR, GCLC, and HEKn, cells were seeded in a 60 mm dish. The day after, cells were treated with derivatives active concentrations. These cytoprotective enzyme levels (ab207621 for HO-1, ab23363 for GCLC, and Cayman Chemical-703202 for GR) were assessed according to the manufacturer’s instructions. Data were given as fold-change compared DMSO-treated cells with SDs and means (n:3).

### Real-Time PCR Analysis

To determine mRNA expression levels, RNA isolations were assessed via Aurum Total RNA Mini Kit (Bio-Rad, USA, 7326820) according to the manufacturer’s instructions, followed by cDNA synthesis performed *via* Iscript cDNA Synthesis Kit (Bio-Rad, USA, 1708891). Then, expression level analysis was conducted via SYBR Green 1 (Bio-Rad, USA, 1725121) and LightCycler480 Thermocycler (Roche, Switzerland). For RT-PCR studies, a specific primer to NRF2 was used (**Supplemental Table S2)**. Transcripts’ fold change was conducted by the normalization with GAPDH as a housekeeping gene. Analysis of Ct values was performed using the Qiagen Rest 2009 program. Experiments were done at three independent experiments with three technical replicates.

