## Supplementary material for "Multi-Targeted Effects of Novel Cycloastragenol Derivatives: Enhancing NRF2, Proteostasis, and Telomerase Pathways with p53 Modulation to Delay Replicative Senescence": Supplemantary materials

Figure S1

Yilmaz et al.

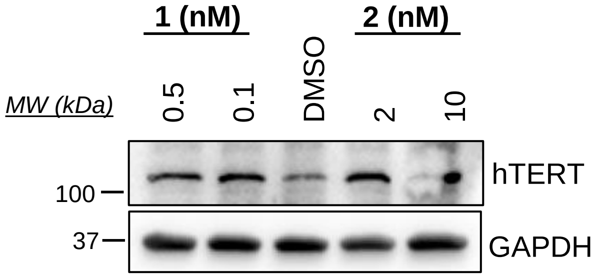


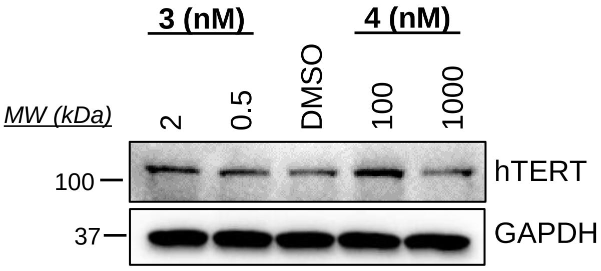


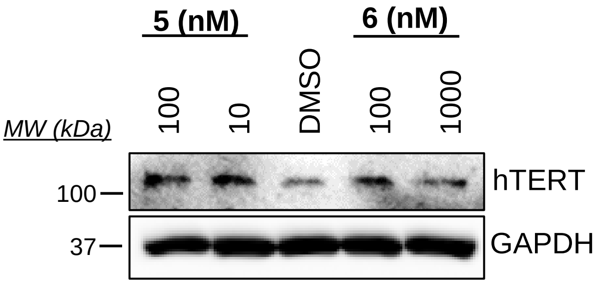

A

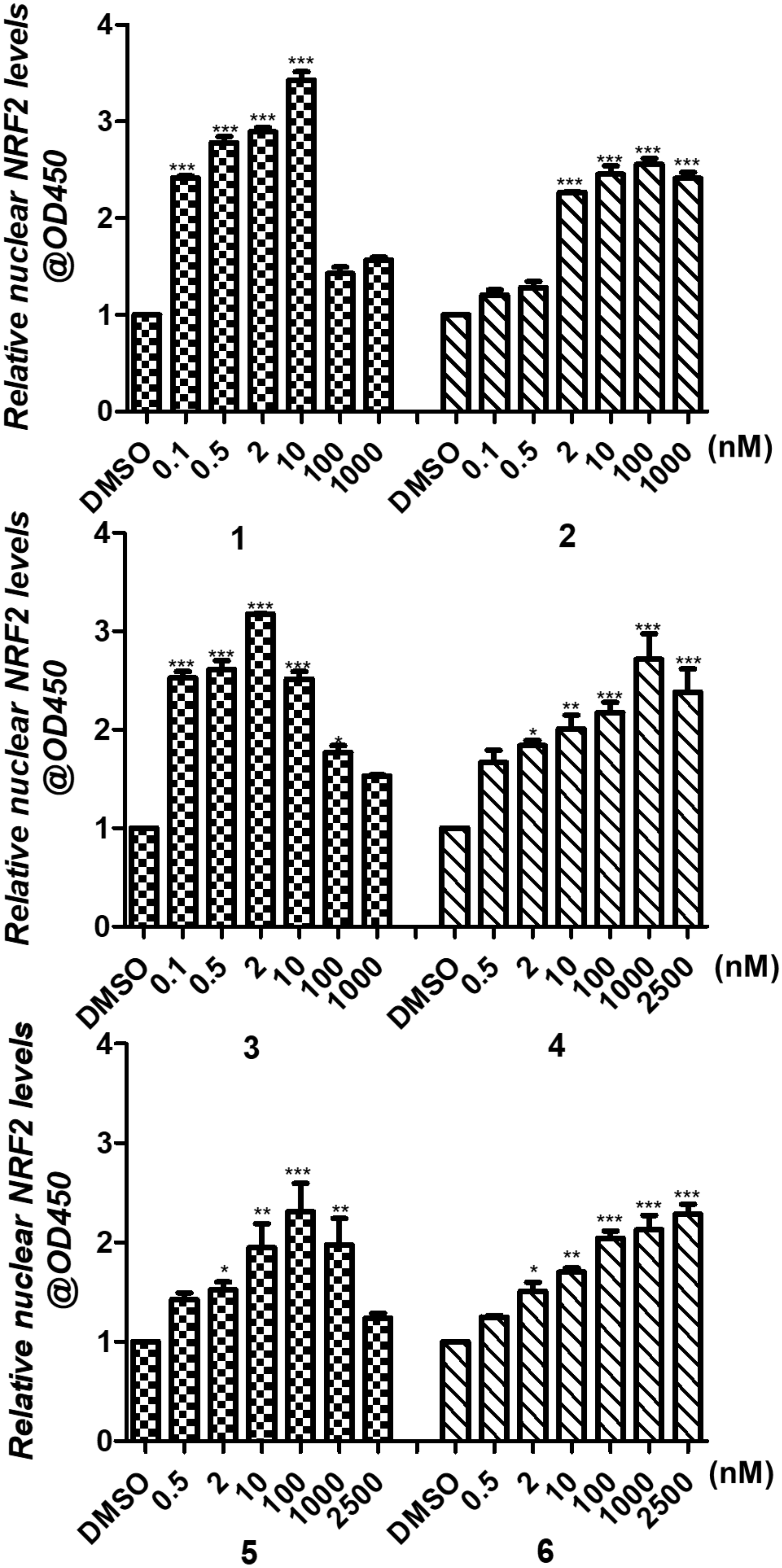
Relative nuclear NRF2 levels @OD450
0
1
2
3
4
***
DMSO
0.1
0.5
2
10
100
1000
(nM)
1
2
3
4
5
6
2500
*
**

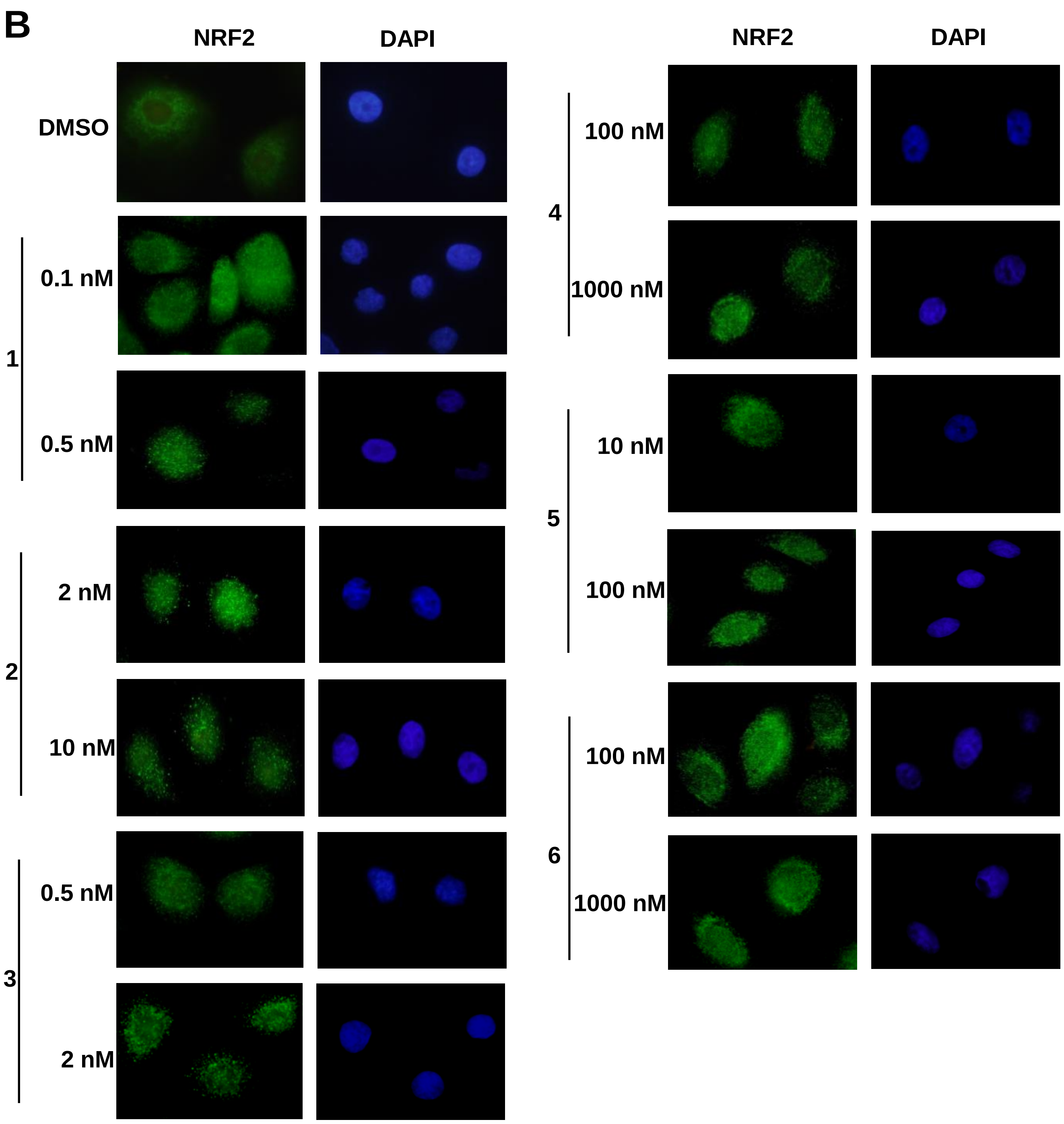
B
NRF2
DAPI
NRF2
DAPI
DMSO
1
0.1 nM
0.5 nM
2
2 nM
10 nM
3
0.5 nM
2 nM
4
100 nM
1000 nM
5
10 nM
100 nM
6
100 nM
1000 nM

**Figure S2**

**Yilmaz et al.**

**C**

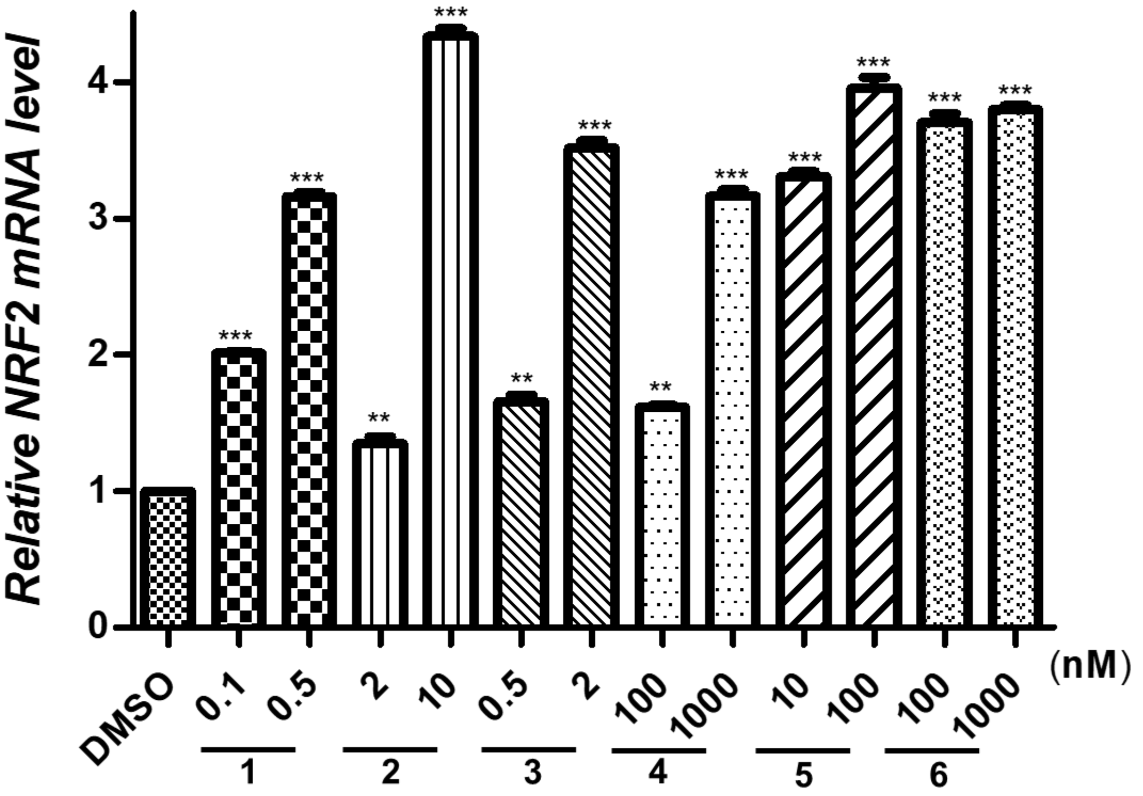


**D**

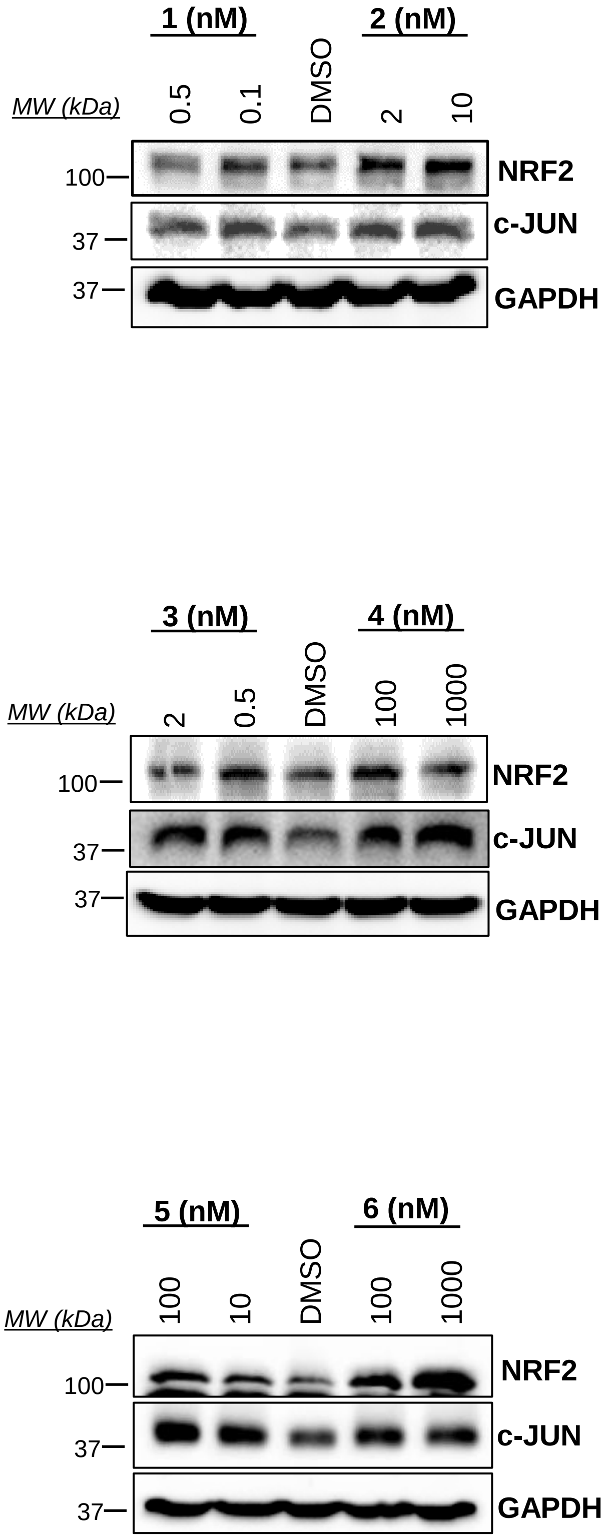

E

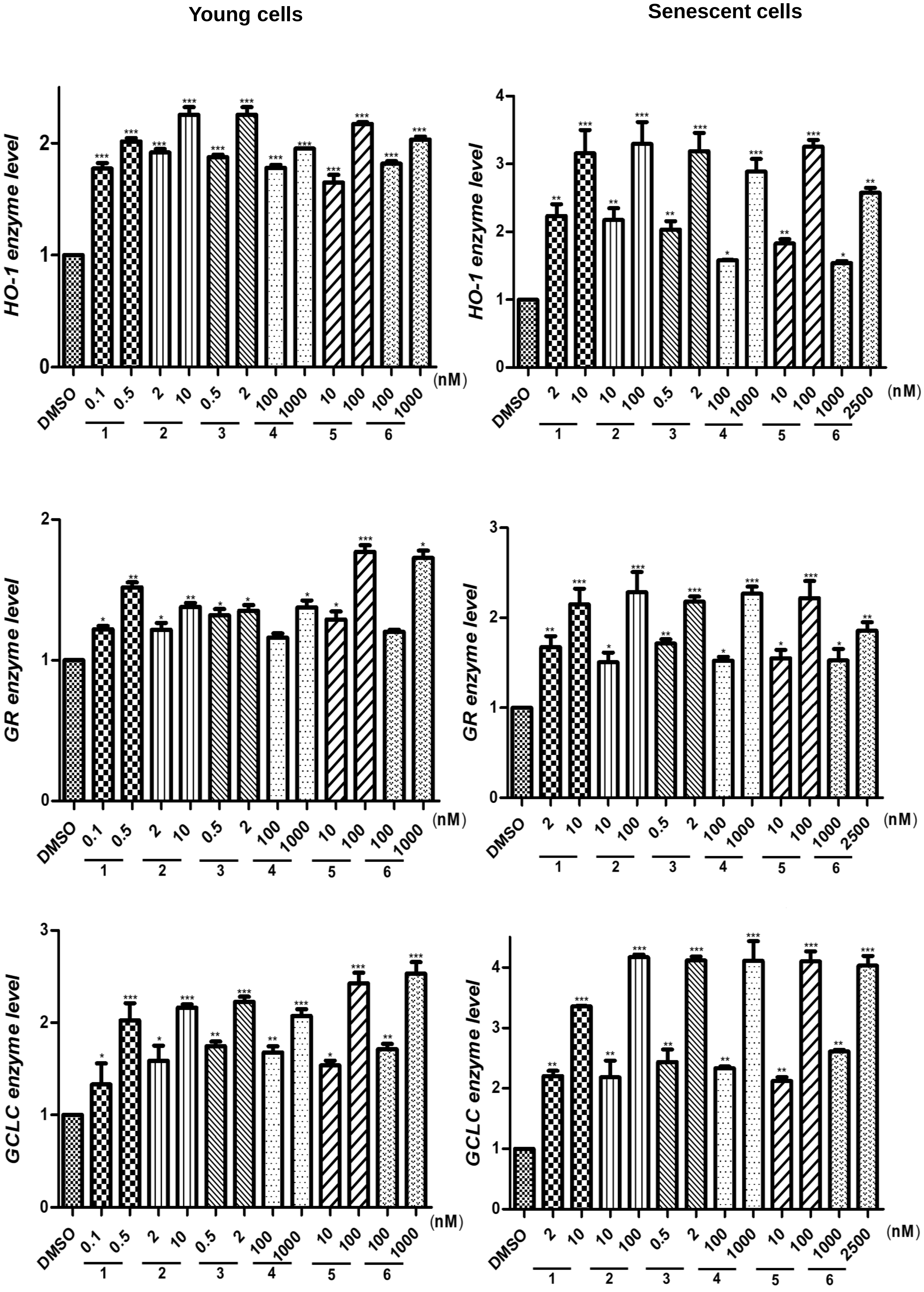
Young cells
Senescent cells
HO-1 enzyme level
GR enzyme level
GCLC enzyme level
DMSO
0.1 0.5 2 10 0.5 2 100 1000 10 100 100 1000
2 10 10 100 0.5 2 100 1000 10 100 1000 2500
(nM)
1 2 3 4 5 6

Figure S3

Yilmaz et al.

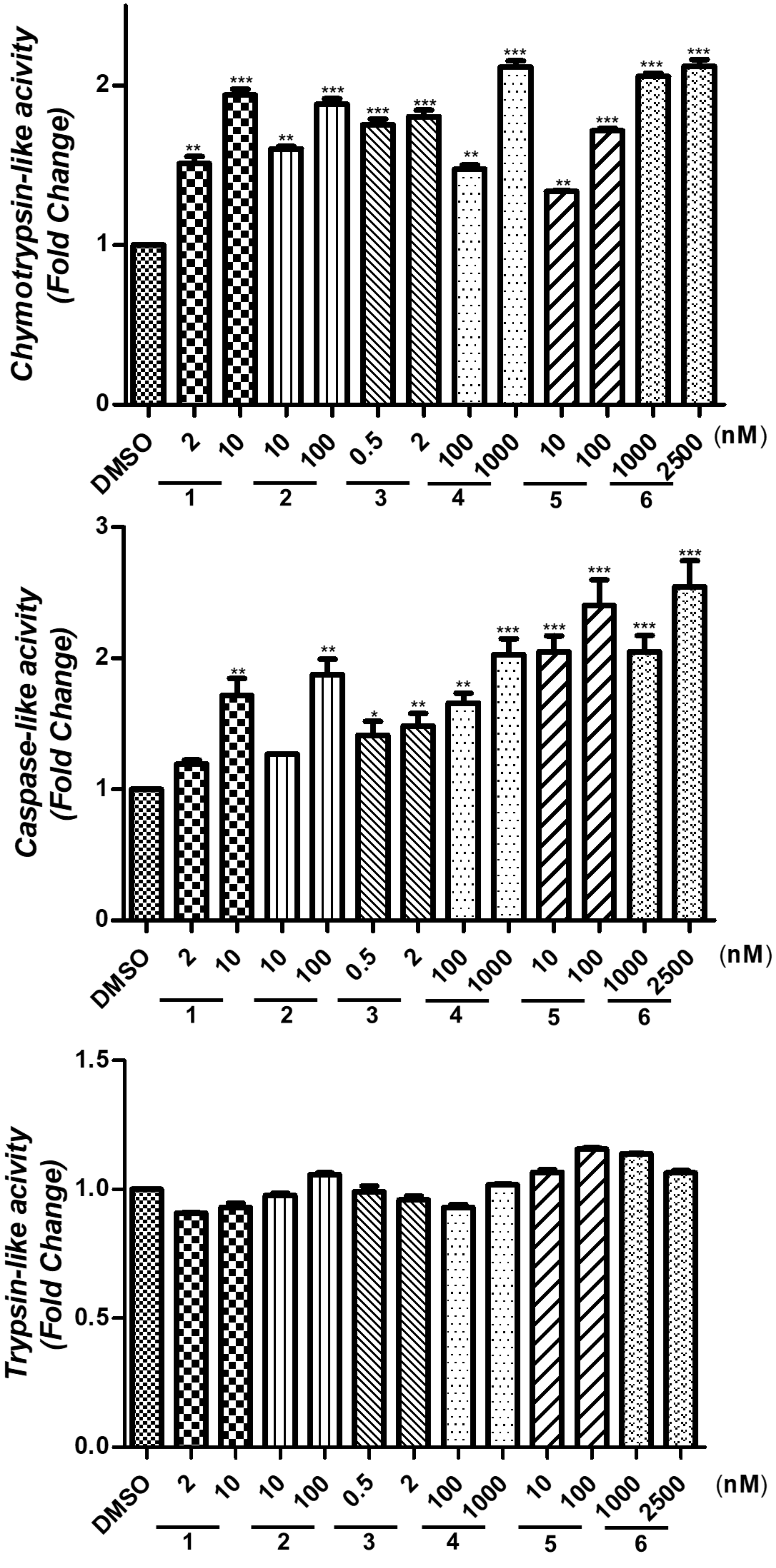

| *Name* | *PDL* |
| --- | --- |
| **DMSO** | 18.78 |
| **CA** | 25.71 |
| **1** | 25.73 |
| **2** | 26.57 |
| **3** | 26.11 |
| **4** | 25.77 |
| **5** | 26.81 |
| **6** | 25.56 |

### Table S2

| Target gene | Forward sequences | Reverse sequences |
|---|---|---|
| NRF2 | GTGAATTTCTCCCAATTCAGCCAG | TCAGGAACAAGTGACTGAAACGTA |

### Figure S4

Yilmaz et al.

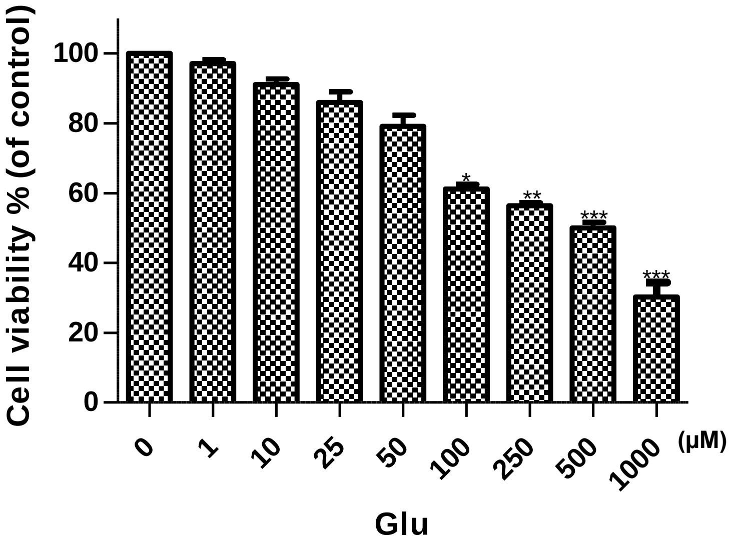

### Figure S5

Yilmaz et al.

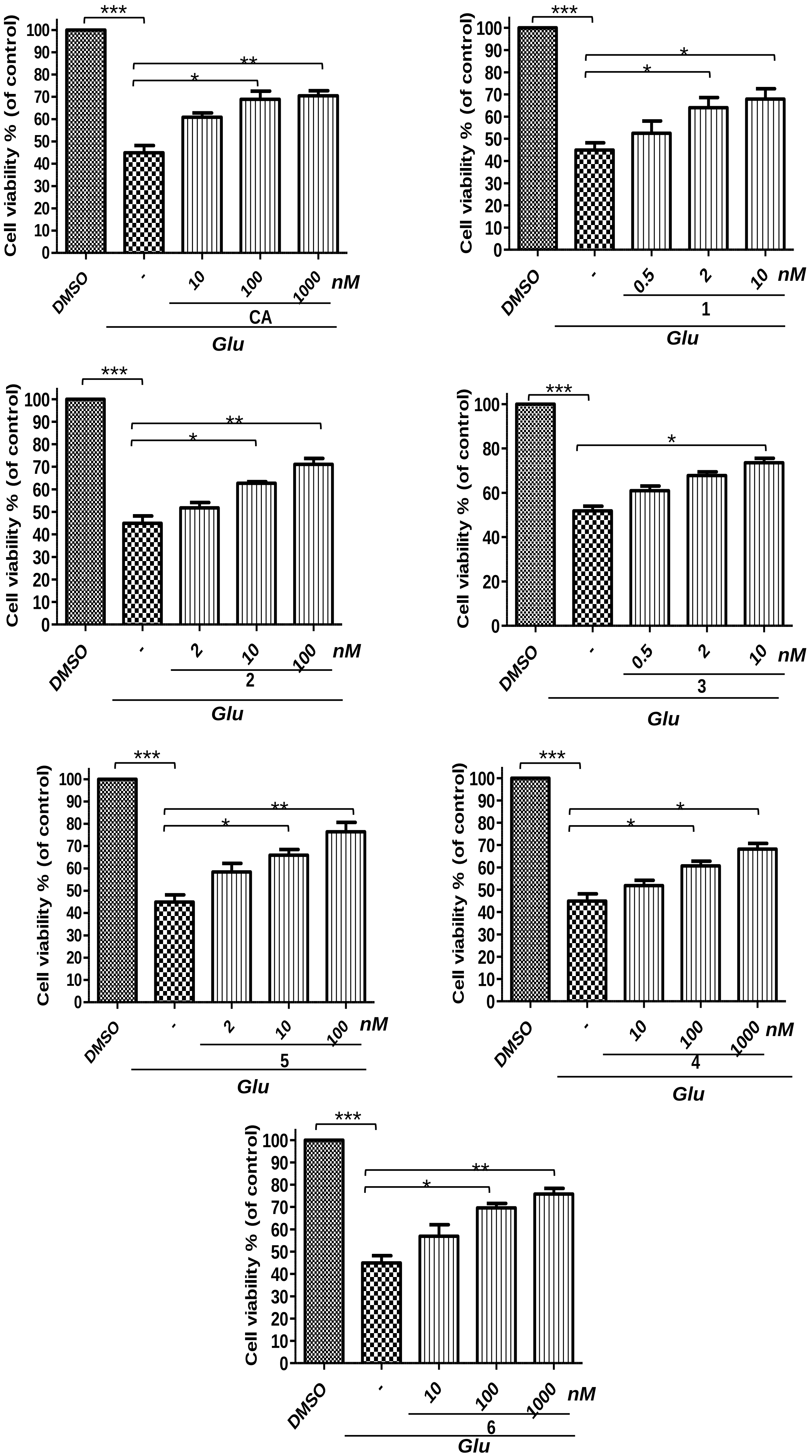
